# Single missense mutations in Vi capsule synthesis genes confer hypervirulence to *Salmonella* Typhi

**DOI:** 10.1101/2023.12.28.573590

**Authors:** Gi Young Lee, Jeongmin Song

## Abstract

Many bacterial pathogens, including the human exclusive pathogen *Salmonella* Typhi, express capsular polysaccharides as a crucial virulence factor. Here, through *S.* Typhi whole genome sequence analyses and functional studies, we found a list of single point mutations that make *S*. Typhi hypervirulent. We discovered a single point mutation in the Vi biosynthesis enzymes that control the length or acetylation of Vi is enough to create different capsule variants of *S.* Typhi. All variant strains are pathogenic, but the hyper-capsule variants are particularly hypervirulent, as demonstrated by the high morbidity and mortality rates observed in infected mice. The hypo-capsule variants have primarily been identified in Africa, whereas the hyper-capsule variants are distributed worldwide. Collectively, these studies increase awareness about the existence of different capsule variants of *S.* Typhi, establish a solid foundation for numerous future studies on *S.* Typhi capsule variants, and offer valuable insights into strategies to combat capsulated bacteria.

## Introduction

Many bacterial pathogens, including the human-exclusive pathogen *Salmonella enterica* serovar Typhi (*S.* Typhi), have the capsular polysaccharides (CPS) biosynthesis system as a crucial virulence factor ^1–8^. Given their role in virulence and the surface display on the cognate pathogen, targeting CPS is most commonly used to prevent those capsulated pathogens ^9–11^. The Vi CPS biosynthesis system of *S.* Typhi is encoded by 9 genes (*tviB*-*E* and *vexA*-*E*) located in the *viaB* locus within *Salmonella* Pathogenicity Island-7 (SPI-7) on the chromosome ^8,12–14^. The expression of the *tviB*-*vexE* operon is regulated by TviA interacting with RcsB or OmpR ^15–17^. TviB and TviC are responsible for synthesizing the Vi monomer (UDP-*N*-acetyl-d-galactosaminuronic acid), which undergoes subsequent polymerization and *O*-acetylation by TviE and TviD, respectively ^8,18,19^. Once the Vi polymer is acylated by VexE ^20^, it is transported to the outer membrane for surface display through the ATP-binding cassette (ABC) transporter composed of VexB-D ^8^.

*S.* Typhi causes the life-threatening systemic disease typhoid fever. Typhoid fever is a major global health concern, as evidenced by outbreaks in Southeast Asia and sub-Saharan Africa ^21–24^. The World Health Organization estimates that the disease claims ∼200,000 deaths per year, mostly children. *S.* Typhi is acquired through ingestion of contaminated food and water, which invades the intestinal mucosa and then spreads systemically to the liver, spleen, bone marrow, and gallbladder ^25^. Following recovery, over 10% of acute typhoid cases become temporary carriers; further, a significant proportion of the individuals infected (2-6%) establish an asymptomatic chronic carriage state (colonization in the gallbladder), during which they excrete *S.* Typhi for months, and in some cases, years ^26–28^. Because *S.* Typhi is a human-specific pathogen, the capacity to generate an asymptomatic carrier state is one of the critical pathogenic mechanisms leading to its persistent presence. Prolonged or persistent infection of *S.* Typhi in the gallbladder and macrophages is known to be a key feature among these persistent asymptomatic carriers ^28–33^. The dynamic infectious cycle of *S.* Typhi is coordinated by many virulence factors, including Vi CPS, flagella, Type III secretion system (T3SS) *Salmonella* Pathogenicity Island (SPI)-1 and SPI-2 effector toxins, and typhoid toxin ^16,29,34–41^.

During the infectious cycle of *S.* Typhi, the Vi capsule serves as a protective barrier against the host’s innate immune responses, exemplified by its roles in inhibiting complement deposition and neutrophil-mediated phagocytosis, while the Vi promotes human macrophage phagocytosis ^42–45^. As a result, it has been observed that acapsular (Vi-negative) *S.* Typhi is generally regarded as avirulent or less virulent in comparison to the wild-type (Vi-positive) *S.* Typhi strains ^42–45^. Nevertheless, there exists a notable deficiency in knowledge regarding the potential emergence of Vi capsule variants and the presence of hypervirulent variants within this category. To address this substantial disparity in knowledge, we conducted bioinformatic analyses, followed by structural and functional investigations. Our findings present experimental evidence that indicates the emergence and circulation of *S.* Typhi capsule variants including hypervirulent Vi variants.

## Results

### *S.* Typhi genes encoding the Vi biosynthesis system are hotspots for clinical missense mutations

The Vi capsule of *S*. Typhi is a crucial virulence factor that distinguishes it from other non-typhoidal *Salmonella* strains. However, despite the significant impact of *S*. Typhi infection on many millions of lives every year, whether any Vi capsule variants have emerged, aside from acapsular strains, remains completely unknown. To fill this fundamental knowledge gap, we first built a bioinformatic pipeline established for the whole genome sequence (WGS) comparison analyses of 5,379 *S.* Typhi clinical isolates (Fig. 1a and Supplementary Table 1). Using this tool, we first conducted a re-annotation of all 5,749 WGSs (available through the NCBI’s GenBank database) using the Prokaryotic Genome Annotation Pipeline (PGAP) to synchronize all annotations under the same criteria. We used RefSeq’s gene numbers to identify the gene pool from each strain and allocated single digit code to each gene: -1, absence of gene, 0, predicted pseudogene, 1, presence of a gene (Fig. 1a and Supplementary Tables 1-2). To ensure that the dataset includes *S.* Typhi WGSs only, we included two additional computational filters in the pipeline: (i) SeqSero2 ^46^ and (ii) typhoid toxin subunits (*cdtB*, *pltA*, and *pltB*) ^47^ to filter out non-Typhi strains, resulting in a total of 5,379 strains that were used in this study (Fig. 1a and Supplementary Table 1).

**Fig. 1.**
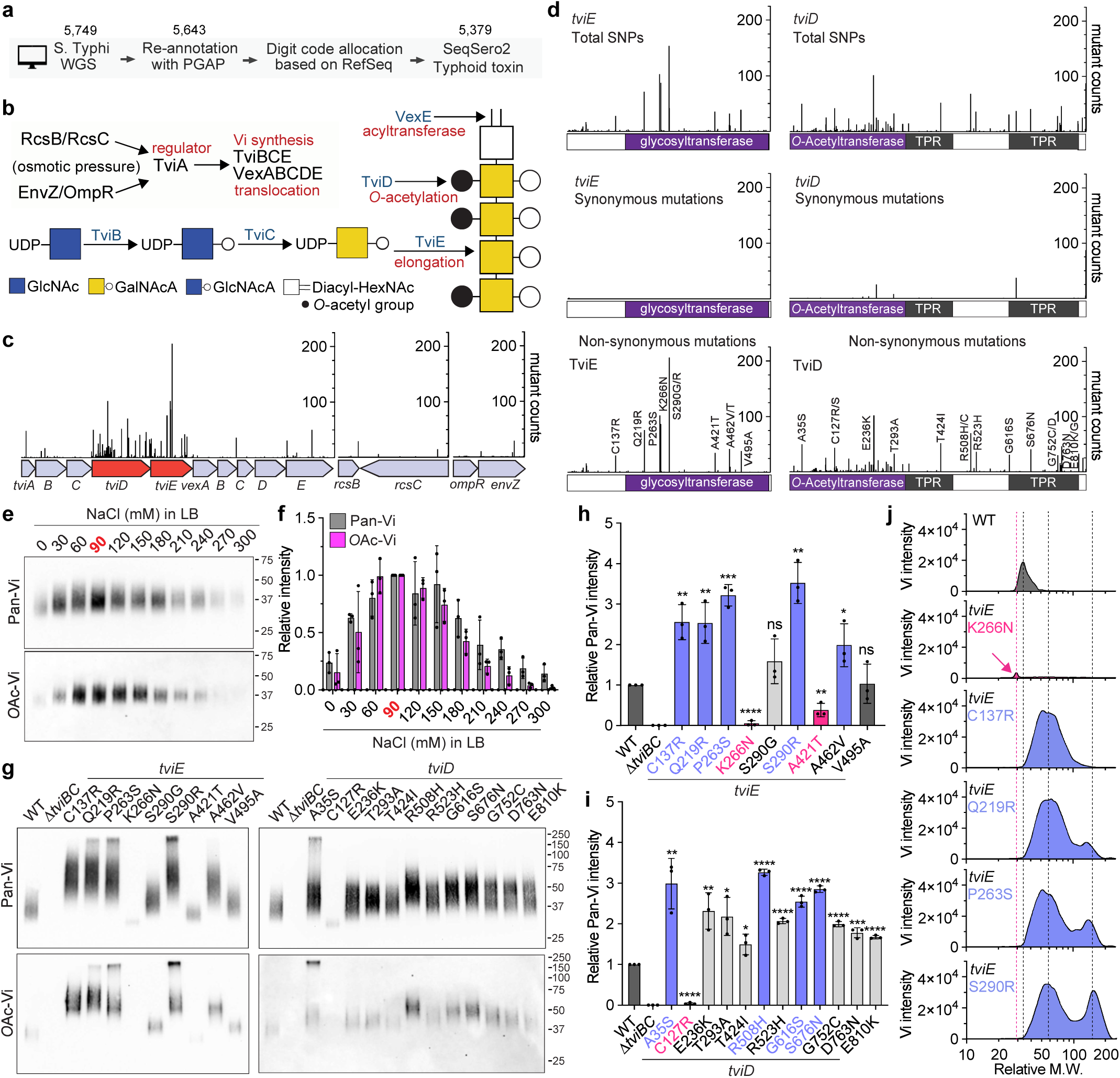
*S.* Typhi genes encoding the Vi biosynthesis system are hotspots for clinical missense mutations. **a,** A schematic workflow of the bioinformatic pipeline used to analyze *S.* Typhi WGSs. **b**, A working model for *S.* Typhi Vi gene regulation, synthesis, modification, and transport. GlcNAc, *N*-acetyl-glucosamine. GalNAcA, *N*-acetyl-galactosaminuronic acid. GlcNAcA, *N*-acetyl-glucosaminuronic acid. Diacyl-HexNAc, diacyl-*N*-acetyl-hexosamine. **c**, Clinical missense mutation frequency map that we have generated in this study. **d**, SNPs (upper panel), synonymous (middle panel), and non-synonymous (lower panel) mutations found in the *tviE* and *tviD* genes. Y-axis, numbers of *S.* Typhi clinical isolates carrying the indicated single point mutation of the indicated genes. X-axis, the position of the single point mutations. TPR, tetratricopeptide repeat. **e**, Immunoblots assessing Vi produced by WT *S.* Typhi cultured in LB containing indicated NaCl concentrations. **f**, Quantification results of three independent experiments associated with e. **g**, Immunoblots assessing Vi produced by *S.* Typhi WT, Δ*tviBC*, and clinical missense mutants cultured in LB containing 86 mM NaCl. **h**, Quantification results of three independent experiments associated with g left panel. **i**, Quantification results of three independent experiments associated with g right panel. **j**, Histograms of Vi length (X-axis) and amount (Y-axis) produced by WT *S.* Typhi and indicated clinical missense mutants. Bars in graphs h-i represent the mean ± standard deviation (SD). Two-tailed student t-tests between WT and indicated mutants were performed. *, P< 0.05. **, P < 0.01. ***, P < 0.001., ****, P < 0.0001. ns, not significant. See also Supplementary Figs. 1-3 and Tables 1-4.

To gain a comprehensive understanding of whether there are mutations in Vi synthesis and its related genes, we analyzed missense point mutations that occurred in the genes for Vi regulation (*rcsB, rcsC, envZ, ompR*, and *tviA*), synthesis (*tviB, tviC*, and *tviE*), modification (*tviD* and *vexE*), and transport (*vexA, vexB, vexC*, and *vexD*) ^8,12,13,18,48,49^ (Fig. 1b). The resulting clinical missense mutation frequency map reveals that nearly all mutations occurred in two genes encoding TviE and TviD (Fig. 1c and Supplementary Table 3).

Contrary to non-synonymous mutations (Fig. 1c), synonymous mutations in the *tviE* gene (1.3%, 10 out of 747 SNPs) and the *tviD* gene (13.7%, 177 out of 1289 SNPs) are much less frequent than non-synonymous mutations (Fig. 1d). In addition, certain strains possess the Vi biosynthesis genes as pseudogenes due to the presence of internal stop and frameshift mutations (Supplementary Table 2). As reported previously ^50,51^, our WGS dataset includes clinical isolates that lack the entire Vi synthesis operon (Supplementary Table 2). *S.* Typhi mutants containing synonymous mutations and predicted pseudogenes are expected to exhibit phenotypic similarities to WT and acapsular *S.* Typhi strains, respectively. Consequently, these strains were excluded from further characterization in this study.

Given their predicted roles within *S.* Typhi in controlling Vi length and/or 3-*O*-acetyl (*O*Ac) modification ^19^, we reasoned that *S.* Typhi variants carrying a single amino acid point mutation in TviE or TviD express a variant form of Vi on the surface of *S.* Typhi. To test the hypothesis, we chose the top 21 clinical missense mutations that are most frequently found in *tviE* and *tviD*, as well as 1 *tviA* mutant. We have generated these 22 mutants of *S.* Typhi, each carrying a single point mutation in *tviE, tviD*, or *tviA*. These mutations were introduced into the isogenic background of the strain ISP2825, which is a clinical isolate of *S.* Typhi ^52^ (Fig. 1d lower panel and Supplementary Table 4).

We first examined Vi production in various salt concentrations (a known environmental cue altering *S.* Typhi Vi expression) ^53^. We observed a higher Vi expression and 3-*O*-acetylation in *S.* Typhi cultured in a medium containing 60-150 mM NaCl with a peak expression at 90 mM NaCl, while a decreased Vi expression was seen in *S.* Typhi cultured in salt-free and high salt-containing media (Fig. 1e-f, and Supplementary Fig. 1). Based on this result, we used 86 mM NaCl LB (a.k.a., low salt LB or Lennox LB) throughout this study, unless specified. Using the validated antibodies specifically detecting pan-Vi (both unmodified and *O*Ac-Vi) or *O*Ac-Vi (Supplementary Fig. 1), we assessed 22 *S.* Typhi mutants and found for the first time that, except *tviA* V62I, each of these single point mutations leads to the overall structural modifications of Vi surface glycan in terms of abundance, length, and *O*-acetyl modification (Fig. 1g-j). Unlike other mutations, we found that *tviA* V62I is not a phenotype-changing mutation (Supplementary Fig. 2). Moreover, our findings indicate that the observed modifications in Vi CPS resulting from individual point mutations are unlikely to be attributed to the altered expressions of TviD or TviE, as we found their comparable mRNA expressions between WT and capsule variant *S.* Typhi strains (Supplementary Fig. 3). Based on the obtained results, the 21 mutations that alter the phenotype were categorized into three distinct groups: hypo-, hyper-, and super-capsule variants. These groups correspond to mutations that result in decreased (pink), highly increased (lilac blue), and intermediately increased (light grey) phenotypic changes, respectively.

### TviE is a processive enzyme, which is altered by clinical missense mutations

To determine the underlying molecular mechanism of how a single amino acid sequence variation alters the Vi length and expression, as well as to identify a plausible molecular evolution strategy used by *S.* Typhi, we predicted the 3-dimensional structure of TviE by running two machine-learning-based structure prediction programs AlphaFold v2 and RosettaFold ^54,55^ (Fig. 2a). We found a strong consensus between the two resulting structures, particularly for the C-terminal and internal regions of TviE, where those clinical missense mutations are positioned (Fig. 2b-c). When we positioned all TviE single point mutations on the 3D structure (which we note that each clinical isolate of *S.* Typhi carries one single point mutation), we noticed that those mutations are not only positioned on the one-themed location, referred to as the “horizontal cleft”, but also result in a drastic change of their electric surface charge distribution of the horizontal cleft (Fig. 2b-c). Based on the aforementioned observations, it can be deduced that amino acids on the horizontal cleft are vital for the catalytic process of the TviE enzyme. More specifically, the long horizontal cleft after the catalytic dyad (E483 and E491)^19^ strongly suggests the “ultra-fast elongation” processive mode in the TviE catalytic process. Regarding this matter, the presence of a single amino acid variation in the cleft is likely to be adequate in altering the interaction dynamics between the enzyme and Vi elongation during action. Consequently, this is likely to be the cause of varying quantities of hypo or hyper Vi produced by these variants (Fig. 2c-d). In accordance with our prediction, we have observed that Vi products produced by WT *tviE* K266, *tviE* K266N (hypo-capsule clinical isolate), *tviE* K266R (positively charged basic a.a.), and *tviE* K266E (acidic a.a.) are in agreement with the proposed TviE catalysis mode (Fig. 2e). This finding provides support for our hypothesized mechanism of adaptive evolution, emphasizing the substantial impact of particular amino acid residues on the elongation process of Vi.

**Fig. 2.**
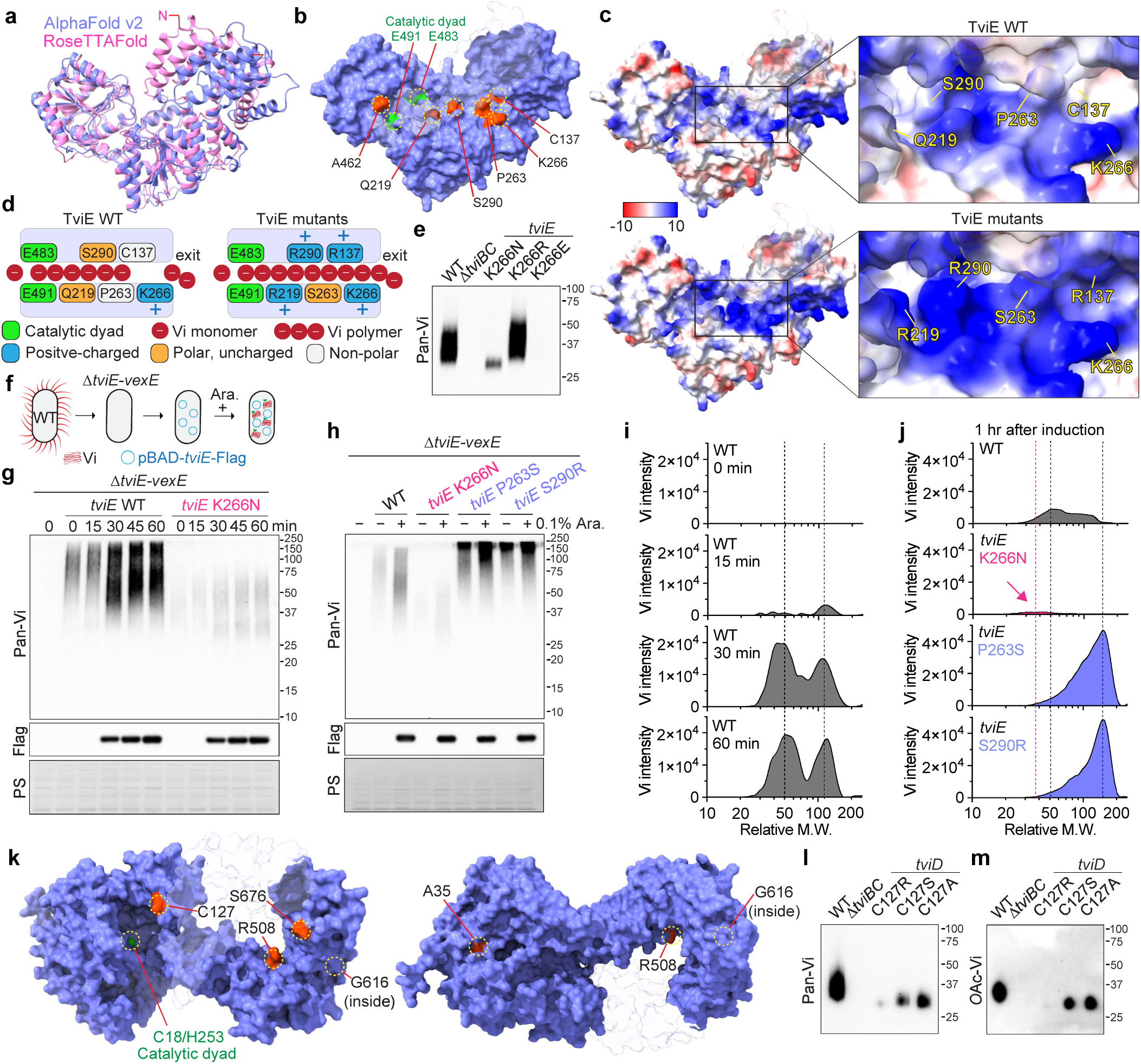
The presence of clinical missense mutations leads to alterations in the mechanism of action of TviE and TviD. **a,** Overlay ribbon diagrams of TviE structures. Pink, TviE structure predicted by RoseTTAFold. Lilac blue, TviE structure predicted by AlphaFold v2. N, N-terminus of TviE. **b**, Clinical missense mutations on the predicted TviE structure (red residues). E491 and E483 are catalytic dyad residues. Lilac blue, the molecular surface of AlphaFold-predicted TviE structure. Light grey/semi-transparent, M1-M45, T214-R218, and R406. **c**, Surface charge distributions of the TviE WT and mutants. Note that R137, R219, R290, and S263 are indicated in the same structure in this figure. Blue, positive charge. Red, negative charge. Semi-transparent, M1-M45, T214-R218, and R406. **d**, A schematic cartoon depicting the predicted Vi extension process occurring on the horizontal cleft of TviE WT and mutants. Red circle, Vi monomer, or polymer. **e**, Immunoblot assessing the role of TviE Lys residue at 266 in Vi length control. *S.* Typhi WT and indicated *tviE* mutants were generated and characterized. **f**, A schematic cartoon illustrating the *in vivo* molecular tool developed to evaluate TviE mutants. **g-h**, Time-course immunoblots of WT *tviE* and *tviE* mutants. PS, Ponceau S stained membranes which was used to demonstrate comparable sample loading. Arabinose (0.1%) was used. **i**, Time-course histograms of Vi length (X-axis) and amount (Y-axis) produced by *S.* Typhi Δ*tviE-vexE* carrying pBAD-*tviE*-Flag. **j**, Histograms of Vi length (X-axis) and amount (Y-axis) produced by *S.* Typhi Δ*tviE-vexE* carrying pBAD-*tviE*-Flag or pBAD-*tviE* K266N, P263S, or S290R-Flag. One hour after 0.1% arabinose induction. **k**, Clinical missense mutations on the predicted TviD structure (red residues). C18 and H253 are catalytic dyad residues. Semi-transparent, the residues A701-S831. **l-m**, Immunoblots assessing the role of TviD Cys residue at 127 in Vi length and *O*-acetylation.

To gain a deeper understanding of the mechanism underlying mutations that confer hypo- and hyper-capsule phenotypes, we generated the *in vivo* molecular tool. We first made a clean-deletion of 6 consecutively located genes within the *viaB* locus in *S.* Typhi, *tviE*, *vexA, vexB, vexC, vexD*, and *vexE*, designated as *S.* Typhi Δ*tviE-vexE* (Fig. 2f). The generated hexadruple mutant was transformed with a plasmid expressing pBAD-TviE-Flag (Fig. 2f). pBAD promoter was used to fine-tune the expression of TviE, while the Flag tag was added to monitor the expression of TviE upon arabinose treatment. Due to the lack of the transporter of the synthesized Vi in this engineered strain (VexA-D), the synthesized Vi is retained within *S.* Typhi, enabling a precise time-course evaluation of the synthesized Vi (Fig. 2f). Using this tool developed, we assessed the impact of 3 representative mutants of TviE, K266N (hypo Vi), P263S (hyper Vi), and S290R (hyper Vi), on Vi synthesis. We found that Lys at position 266 is critical for the mode of action for TviE, as K266N resulted in hypo Vi product with a reduced expression (Fig. 2g). In contrast, we found that the mutations associated with the hyper-capsule Vi product (P263S and S290R) resulted in a stronger expression of hyper Vi than wild-type (WT or predominant strain) *S.* Typhi (Fig. 2h-j), indicating that Pro at position 263 and Ser at position 290 are also critical for the catalytic mode of TviE. Intriguingly, all critical residues identified in this study are located in the internal region of TviE, which serves as the first example of its kind across bacterial glycosyltransferases involved in capsule synthesis.

### Clinical missense mutations of TviD are distributed throughout its 3D structure, yet it seems to evolve to acquire specific amino acid residues for specific variants

To gain a mechanistic understanding of the effects of TviD’s clinical missense mutations on Vi production, we have employed the predicted structure of TviD to superimpose the frequently observed clinical missense mutations (Fig. 2k). In contrast to the frequent clinical missense mutations observed in TviE, the mutations in TviD are distributed throughout its 3D structure, specifically on its surface except G616. The surface and distributed location of these mutations suggest that multiple adaptive evolutionary events have contributed to their emergence. These mutations are likely to have impacts on various aspects of Vi synthesis process, including TviD’s catalytic activity and the formation of the predicted multimeric, multiprotein enzyme complexes for Vi production. Further investigation is required to comprehend the specific effects of each mutant on Vi production. However, for the purpose of this study, we have chosen to focus on the C127R variant (hypo-capsule). This particular variant is situated in close proximity to the catalytic site of TviD, approximately 17 Å and 21 Å away from H253 and C18, respectively (Fig. 2k). Our aim was to experimentally assess the influence of specific amino acid residues, as observed in clinical isolates, on Vi production (Fig. 1g). We found that the clinical missense mutation C127R significantly impairs the function of TviD (Fig. 2l-m). In contrast, experimental mutations, specifically C127S and C127A, result in a modest decrease (Fig. 2l-m). These findings offer evidence in favor of bacterial adaptive evolution, as the substitution of residue C127 with residue R appears to exhibit a high degree of specificity. Based on the close distance to the catalytic dyad, it can be inferred that the C127 residue is potentially involved in facilitating the transfer of the acetyl group to Vi during the acetylation process of TviD.

### *S.* Typhi hypo-capsule variants possess several critical pathogenic characteristics, including serum resistance and increased invasion

We have observed similar growth patterns in both the *S.* Typhi WT and Vi variant strains (Fig. 3a). This suggests that there is no discernible fitness cost associated with the production of variant forms of the Vi capsule. It is equally important to ascertain whether Vi variants, similar to WT *S.* Typhi, possess pathogenic properties. We reasoned that these hypo- and hyper-capsule Vi variants are pathogenic since these clinical mutations are originated from human infection sites (Fig. 3b and Supplementary Table 1). In particular, in the so-called “hypo-capsule” group (>3% of 5,379 clinical isolates; Fig. 3b), *S.* Typhi hypo Vi variants exemplified by *tviE* K266N and *tviD* C127R have thin Vi capsule, which, therefore, we hypothesized that the hypo-capsule variants are able to avoid host innate immune defense mechanisms like WT (e.g., serum resistance), while these variants invade host cells better than WT. In the “hyper-capsule” group (>8%; Fig. 3b), *S.* Typhi hyper Vi variants (e.g., *tviE* C137R, *tviE* Q219R, *tviE* P263S, and *tviE* S290R) are expected to demonstrate an increased capacity to evade neutrophil-mediated phagocytosis and other immune responses. We predict that this is attributed to the presence of a thick Vi capsule and the shed Vi antigen. The “super-capsule” group (>8%: Fig. 3b) is anticipated to have an intermediate phenotype between WT and hyper-capsule variants.

**Fig. 3.**
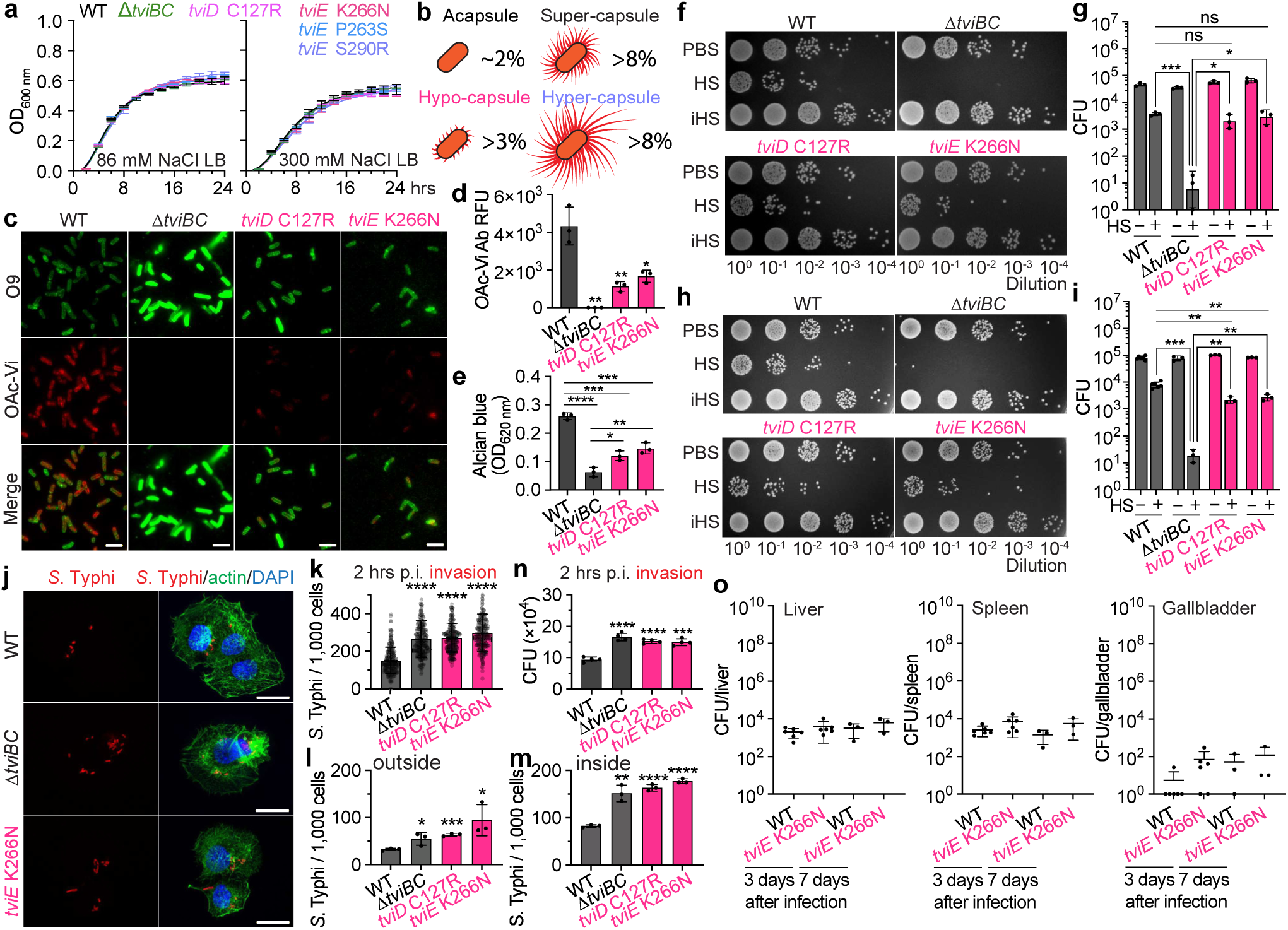
*S.* Typhi hypo-capsule variants possess several critical pathogenic characteristics, including serum resistance and increased invasion. **a,** Growth curves of *S.* Typhi WT, Δ*tviBC*, and indicated clinical missense mutants cultured in 86 mM NaCl LB (Lennox LB) or 300 mM NaCl LB. Lines represent the mean ± SD. **b**, Schematic illustration depicting the capsule variants. Numbers in % indicate the % of capsule variants found among 5,379 clinical isolates. **c**, Representative images of image cytometry analyses on *S.* Typhi WT, Δ*tviBC*, *tviD* C127R, and *tviE* K266N. scale bars, 5 µm. **d**, Quantification results of three independent experiments associated with c. **e**, Alcian blue staining results of three independent experiments. **f-i**, Human serum resistance assays of *S.* Typhi WT and mutants. Ten-fold dilutions of bacterial cultures, which were incubated at 37°C for 2 hours in the presence of 90% non-vaccinated **(f-g)** or vaccinated **(h-i)** human sera, plated on LB agar plates, incubated overnight, and photographed. PBS, no sera. HS, human sera. iHS, heat-inactivated human sera. **j**, Representative fluorescence microscopy images of bacterial invasion into host cells. Henle-407 cells were infected for 2 hours with 15 m.o.i. of *S.* Typhi WT, *ΔtviBC*, *tviE* P263S, or *tviE* K266N. Scale bars, 20 µm. **k**, Quantification results of three independent experiments associated with j. 2 hours after infection. **l-m**, Quantification results of three independent experiments assessing outside **(l)** and inside **(m)** *S.* Typhi 1 hour after infection. **n**, Quantification results of three independent experiments assessing CFUs of intracellular *S.* Typhi 2 hours after infection. Bars represent the mean ± SD except graph g and I that show the geometric mean ± SD. Two-tailed t-tests between WT and indicated strain were performed unless otherwise indicated. *, P< 0.05. **, P < 0.01. ***, P < 0.001., ****, P < 0.0001. ns, not significant. **o**, CFU assays in the liver, spleen, and gallbladder 3 and 7 days after infection. Mann-Whitney tests were performed.

To demonstrate the thin capsule of hypo-capsule variants, we conducted image cytometry using antibodies specific to Vi and LPS O9 surface glycans (Fig. 3c). We indeed observed that the hypo-capsule variants *tviE* K266N and *tviD* C127R of *S.* Typhi exhibit minimal Vi expression on the bacterial surface (Fig. 3c-d). This level of expression is significantly lower than that of the WT bacteria, but differs from the complete absence of Vi expression observed in the acapsular *S.* Typhi *ΔtviBC* strain (Fig. 3c-d). To conduct a further assessment of the limited presence of Vi on the surface of hypo-capsule variants, we employed alcian blue 8GX. This particular stain is widely used for detecting acidic, neutral, sulfated, and phosphate polysaccharides as well as glycosaminoglycans ^56^. In accordance with the image cytometry results, we observed a significant decrease in alcian blue staining on *S.* Typhi with the mutation that confers hypo-capsule formation, compared to the WT strain (Fig. 3e). However, the staining intensity was significantly higher than the acapsular strain that only expresses LPS O9 glycan (Fig. 3e).

We conducted an investigation to assess the serum resistance of the hypo-capsule variants. This was accomplished by utilizing human sera obtained from unvaccinated healthy individuals who had not been immunized against typhoid fever. In contrast to the acapsular *S.* Typhi *ΔtviBC* strain, the survival of *S.* Typhi strains carrying *tviD* C127R and *tviE* K266N, which are two hypo-capsule variants, remained robust even after 2 hours of incubation at 37°C in the presence of 90% healthy human sera. These strains exhibited only a minor decrease in survival when compared to the WT *S.* Typhi strain (Fig. 3f-g). The specificity of the findings (Fig. 3f-g) is supported by the use of PBS-only or heat-inactive human sera (iHS) samples. The findings suggest that even a low level of Vi expression on *S.* Typhi is effective in conferring resistance to serum, thereby distinguishing the hypo-capsule Vi variants from the acapsular counterparts. Comparable results of serum resistance of the hypo-capsule variants were also observed with human sera from vaccinated healthy individuals who had been immunized against typhoid fever (Fig. 3h-i).

One significant pathogenic characteristic that sets *S.* Typhi apart from other bacterial pathogens with capsules is its superior ability to invade host cells. Our research, conducted using quantitative microscopy-based infection assays, revealed that hypo-capsule variants significantly enhanced the infectivity of *S.* Typhi by approximately 300% (Fig. 3j-k). This suggests that *S.* Typhi Vi variants with hypo Vi exhibit an expanded pathogenic characteristic. By employing quantitative inside-outside staining, we have additionally discovered that hypo-capsule Vi variants of *S.* Typhi exhibit a higher degree of adherence to host cells compared to the WT strain (Fig. 3l-m), which is consistent with their notable increase in invasion. These results are in agreement with the findings from bacterial colony forming unit (CFU) assays (Fig. 3n). Next, we proceeded to assess whether the hypo-capsule variant demonstrates an extended invasion persistence using Cmah null mice. The Cmah null mice have an intact murine immune system while exclusively expressing glycans that are terminated in N-acetylneuraminic acid (Neu5Ac) due to the lack of the CMP-Neu5Ac hydroxylase (Cmah), which closely resemble those observed in humans ^57^. In contrast, the C57BL/6 mice (having the functional Cmah enzyme) exhibit glycans that are terminated in both Neu5AC and N-glycolylneuraminic acid (Neu5Gc). It is important to acknowledge that the brain of C57BL/6 mice, like that of humans and Cmah null mice, exclusively expresses Neu5Ac, highlighting that the expression of Cmah varies across different cells and tissues in mice ^58^. Due to a closer glycan resemblance of Cmah null mice to humans, these animals have been used for studying the virulence of human pathogens ^47,59–64^. We observed that mice infected with WT and the hypo-capsule variant were comparable, although organ CFUs of the hypo-capsule variant were modestly increased, compared to that of WT (Fig. 3o). The results suggest that the hypo-capsule variants may exhibit a longer persistence in infected individuals.

### The hyper-capsule variants of *S.* Typhi are hypervirulent with both bacteria-associated and shed Vi contributing to their heightened virulence

To assess the pathogenicity of the hyper-capsule variants *in vivo*, we infected Cmah null mice with WT or hyper-capsule (*tviE* P263S) *S.* Typhi. Mice infected intraperitoneally with 8 x 10^5^ of the hyper-capsule variant (*tviE* P263S) showed severe clinical signs (including diarrhea, overall weakness, and eye discharge) and liver abscesses, resulting in the death of 50% of the infected mice by day 3 post infection (Fig. 4a-c, and Supplementary Table 5). Organ CFUs of the hyper-capsule variant were markedly increased, which was 100,000-fold higher than that of WT at day 3 post infection (Fig. 4d). It is notable that 100% of mice infected with the hyper-capsule *S.* Typhi successfully colonized in the gallbladder (Fig. 4d). Intriguingly, consistent with the observed superior gallbladder colonization of the hyper-capsule *S.* Typhi, a recent WGS analysis study on human gallbladder carriage *S.* Typhi isolates found other hyper-capsule mutations, *tviD* R508H, *tviE* C137R, and *tviE* A462V in several gallbladder carriage isolates ^33,65^.

**Fig. 4.**
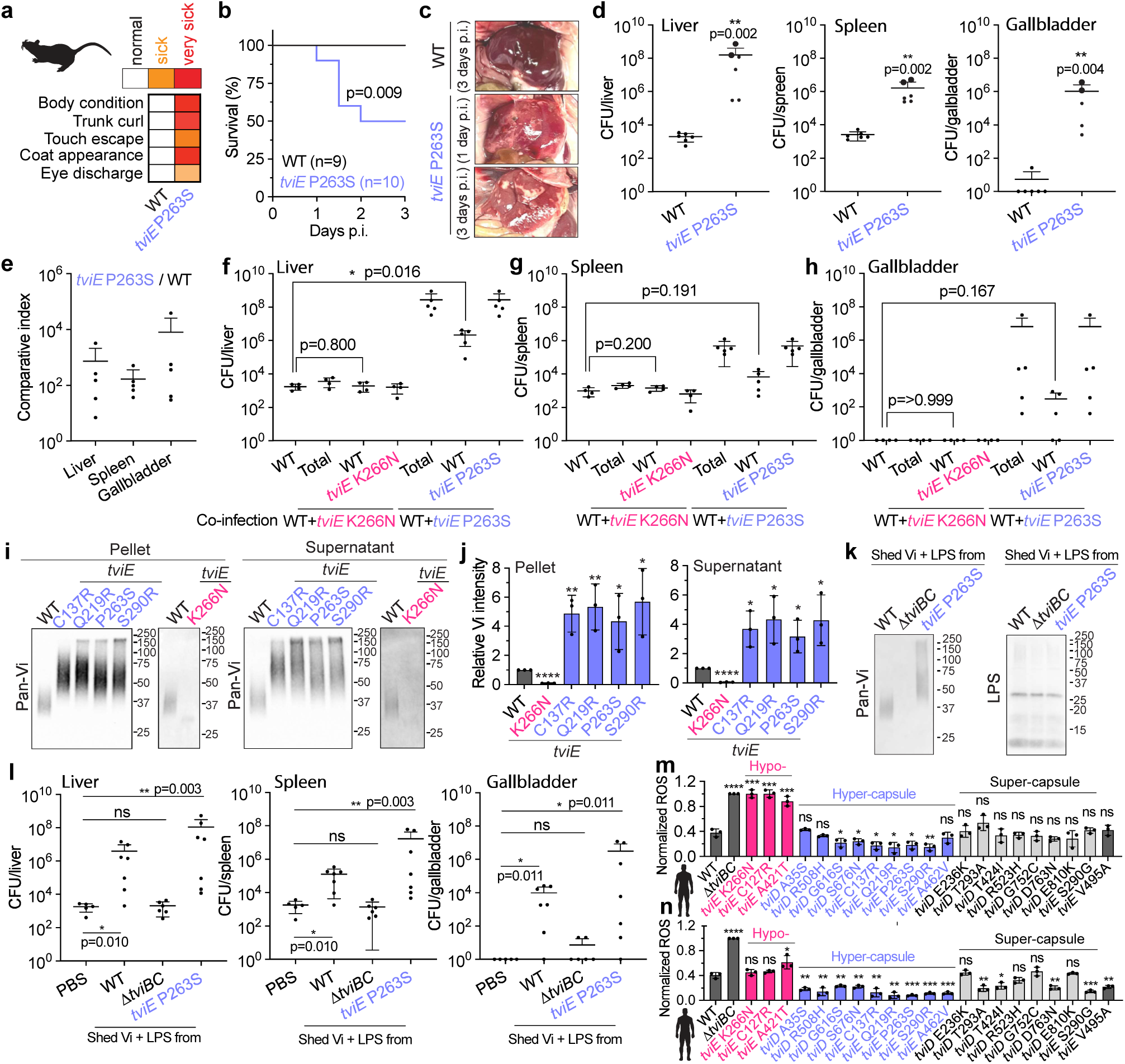
The hyper-capsule variants of *S.* Typhi are hypervirulent, with both bacteria-associated and shed Vi contributing to their heightened virulence. **a-d**, SHIRPA test **(a)**, percent survival **(b)**, and liver abscess **(c)** from Cmah-null mice infected intraperitoneally with 8 x 10^5^ WT (n=9), or *tviE* P263S (n=10). **d**, CFU assays in the liver, spleen, and gallbladder 3 days after infection. Big dots represent CFU numbers from two severely sick mice that died on day 1.5. Note that the organ CFUs of WT *S.* Typhi on day 3 are the same as those of Fig. 3o WT *S.* Typhi on day 3 because these studies were carried out concurrently. **e-h**, Chloramphenicol-resistant or Kanymycin-resistant genes were inserted downstream of the pseudonized *pagL* gene of WT (CmR), *tviE* K266N (KanR), or *tviE* P263S (KanR). Cmah-null mice were infected intraperitoneally with 4 x 10^5^ WT *S.* Typhi, or with a combination of WT+*tviE* K266N (4 x 10^5^ each), or WT+*tviE* P263S *S.* Typhi strains (4 x 10^5^ each). Comparative index **(e)** and CFUs in the liver **(f)**, spleen **(g)**, and gallbladder **(h)** 3 days after infection. **i**, Immunoblots assessing the shedding of Vi by *S.* Typhi *tviE* K266N, *tviE* C137R, *tviE* Q219R, *tviE* P263S, and *tviE* S290R. **j**, Quantification results of three independent experiments associated with i. **k-l**, *In vivo* effects of shed Vi on bacterial infection. Immunoblot analysis of Vi preparations and residual LPS **(k)**. Shed Vi (1.6 x 10^9^ bacteria) was prepped in 100 µL PBS with 8 x 10^5^ WT *S.* Typhi for infection. CFUs in the liver, spleen, and gallbladder 24 hours after infection **(l)**. **m-n**, ROS burst from neutrophils incubated with the indicated strain after opsonization with un-vaccinated **(m)** or vaccinated **(n)** human sera. RLU, relative luminescence unit. Bars/lines represent the mean ± SD. Log rank tests (b), two-tailed t-tests (j) and Mann-Whitney tests (all others) were used to compare WT to the designated strain. *, P< 0.05. **, P < 0.01. ***, P < 0.001., ****, P < 0.0001. ns, not significant. See also Supplementary Table 5.

To further evaluate the impact of the hyper Vi capsule variant on *in vivo* infection of *S.* Typhi, we conducted co-infection studies in mice. The co-infection involved the simultaneous administration of WT and hyper-capsule *S.* Typhi strains at a 1:1 ratio (4 x 10^5^ each). The total inoculum used for the co-infection was equivalent to the single infection inoculum described in Fig. 4a-d. We found that the organ CFUs of hyper-capsule *S.* Typhi were consistently 2-4 log higher than WT (Fig. 4e). Intriguingly, there was an observed increase in the infection of co-infected WT *S.* Typhi (Fig. 4f-h). The phenotype was not observed when WT *S.* Typhi was co-infected with the hypo-capsule variant of *S.* Typhi (Fig. 4f-h), indicating that the observed phenotype is a result of the heightened expression of Vi on hyper-capsule variants.

Based on the findings derived from co-infection studies, it can be inferred that the excretion of Vi antigen by hyper-capsule bacteria potentially is vital for the heightened infection observed in co-infected WT *S.* Typhi. We conducted an assessment to determine if the hyper Vi capsule variant exhibits substantial Vi shedding. Our findings revealed that *S.* Typhi strains with hyper-capsule shed significantly higher quantities of the hyper-capsule Vi (Fig. 4i-j). On the other hand, the shedding of Vi was found to be negligible in the case of hypo-capsule *S.* Typhi (Fig. 4i-j). To investigate the impact of shed Vi on *S.* Typhi infection, we conducted a purification process on the shed Vi. This involved clearing Vi by filtration, removing the majority of LPS, and subsequently purifying it using size exclusion chromatography. The purified Vi (with low LPS contamination) was utilized for conducting infection studies in mice (Fig. 4k). The mice were subjected to co-injection with purified Vi and WT *S.* Typhi strain (Fig. 4l). Preparations of Vi antigen derived from both WT and hyper-capsule variants of *S.* Typhi were utilized. To eliminate the possibility of LPS-mediated effects, we have also included the equivalent preparation from acapsular *S.* Typhi *ΔtviBC*. We observed that the inclusion of Vi preparations significantly increased the infection of WT *S.* Typhi when compared to the absence of additional Vi (Fig. 4l). We also observed that the Vi preparation obtained from hyper-capsule *S.* Typhi tends to exhibit a stronger increase effect than the Vi preparation derived from WT *S.* Typhi (Fig. 4l). The results collectively indicate that the hyper-capsule Vi variant shed a higher quantity of Vi, which is crucial for the observed enhancement of *in vivo* infection in both the hyper-capsule single infection and the co-infected WT *S.* Typhi (Fig. 4d-h).

To understand the role of bacteria-associated hyper-capsule Vi, as well as to gain insights into their relevance to human infection, human peripheral blood neutrophils were isolated from healthy blood donors and assessed neutrophil reactive oxygen species (ROS) burst ^36,66^. *S.* Typhi strains were opsonized with 10% human sera obtained from unvaccinated healthy individuals who had not been immunized against typhoid fever. We found that ROS burst from neutrophils incubated with hyper-capsule variants were much less susceptible to phagocytosis (Fig. 4m). Super-capsule variants had intermediate phenotypes, as some phenocopied hyper-capsule variants while some phenocopied WT (Fig. 4m). Comparable results were also observed with human sera from vaccinated healthy individuals who had been immunized against typhoid fever (Fig. 4n). This study collectively illustrates that hyper-capsule *S.* Typhi is a viable, hypervirulent pathogen possessing specific enhanced pathogenic traits. These characteristics may potentially benefit other Vi variants and even other pathogens in cases of co-infection.

### The hypo-capsule variants have primarily been identified in Africa, whereas the hyper-capsule variants are distributed worldwide

To gain insights into the occurrence, distribution, and persistence of hypo- and hyper-capsule *S.* Typhi variants in human populations, along with their potential association with antibiotic resistance, we conducted comprehensive analyses on clinical isolates of *S.* Typhi. Our analysis included an initial dataset of 5,379 isolates (Fig. 1a), as well as 13,081 WGSs obtained from PathogenWatch. It has been observed in this study that the hypo and hyper-capsule variants, which were characterized, have been persistently present in human populations throughout the entire century (Fig. 5a-d). This indicates the enduring presence of both hypo- and hyper-capsule Vi variants.

**Fig. 5.**
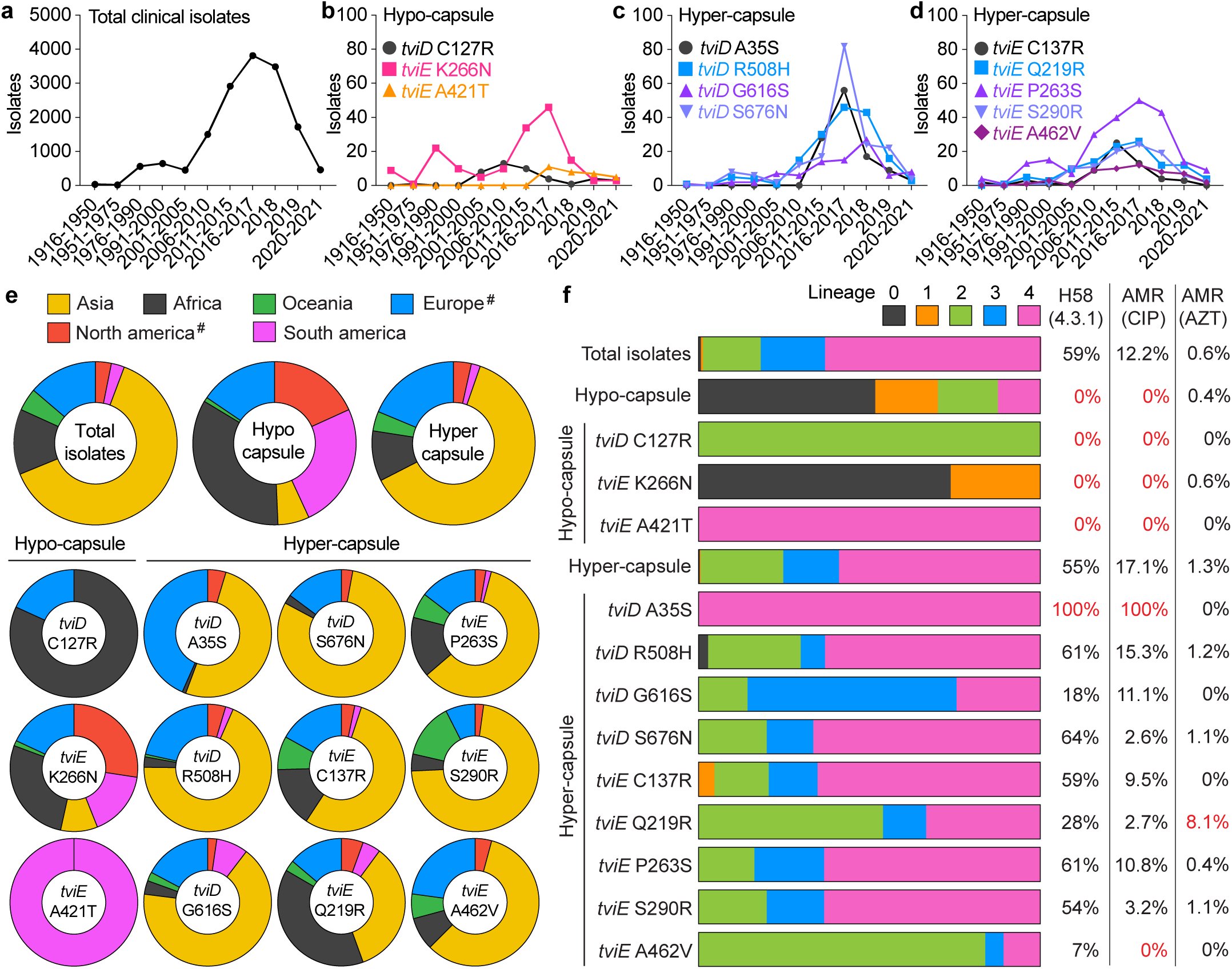
Global distribution of hypo- and hyper-capsule variants. *S.* Typhi WGSs from NCBI and Pathogenwatch were merged and evaluated. Duplicate samples were eliminated, yielding a total of 16,368 strains. **a-d**, The years in which *S.* Typhi Vi variants have been observed. Total (**a**), hypo-capsule (**b**), and hyper-capsule (**c-d**) clinical isolates isolated from 1916 to 2021. **e,** Global distribution of the capsular variants. #, the isolation of *S.* Typhi from North America (USA and Canada) and Europe (UK) may be related to travel. **f**, Capsular variant genotypes and related antimicrobial resistance. See also Supplementary Table 6. CIP, ciprofloxacin. AZT, azithromycin.

Based on our analyses, it has been determined that the hypo-capsule variants carrying *tviD* C127R and *tviE* K266N have been prevalent in various African countries for the past century (Fig. 5e and Supplementary Table 6). In contrast, the hypo-capsule variant carrying *tviE* A421T has emerged in 2017, which has been observed in Brazil (Supplementary Table 6). Consistent with their dissemination findings, the hypo-capsule variants appear to undergo convergent evolution, as the two previous variants of hypo-capsules are classified as *S.* Typhi lineage 0, 1, and 2, while the new variant *tviE* A421T falls under lineage 4 (Fig. 5f). Our analysis did not observe any correlation between hypo-capsule variants and antibiotic resistance (Fig. 5f). The existence of hypo-capsule variants over the past century, despite their susceptibility to drugs, aligns with the mild yet prolonged invasion we observed in a mouse model.

In contrast to the limited distribution of hypo-capsule variants, our analysis indicates that hyper-capsule variants are widely distributed on a global scale (Fig. 5e and Supplementary Table 6). Hyper-capsule variants have also been observed persistently for the entire century, but their distribution is limited to lineages 2, 3, and 4, with the majority being found in lineage 4 (Fig. 5f). Notably, the hyper-capsule *tviD* A35S was first identified in 2013 and has predominantly been observed in India (Fig. 5e and Supplementary Table 6). This particular variant is exclusively found within the lineage 4.3.1.2.1, which is known to be one of the ciprofloxacin-resistant H58 (4.3.1) lineages (Fig. 5f). The findings collectively indicate that there were convergent occurrences of hypo- and hyper-capsule variations. Moreover, all instances of the hyper-capsule *tviD* A35S variant exhibit resistance to ciprofloxacin (Fig. 5f), which has traditionally been the mainstay treatment for patients with typhoid in clinical settings, which therefore suggests the potential for further dissemination of specific hyper-capsule variants.

## Discussion

Here, we performed pan-genome analyses of 5,379 *S.* Typhi clinical isolates to determine whether the capsule variants of *S*. Typhi have emerged. Using the frequent clinical missense mutations in the *viaB* locus identified in this study, we conducted structural and functional studies to understand their consequences in *S.* Typhi pathogenicity and explain the molecular mechanisms involved. Resultantly, we generate a list of single missense mutations found in Vi capsule biosynthesis enzymes that are responsible for producing hypo- and hyper-capsule variants of *S.* Typhi. This paper is the first to demonstrate the functional consequences of clinical missense mutations in the *S.* Typhi *viaB* locus. It serves to highlight the existence of hypo-capsule and hyper-capsule, hypervirulent variants of *S.* Typhi, thereby increasing awareness in the community. These findings provide essential groundwork for enhancing preparedness in response to the potential future spread of hyper-capsule *S.* Typhi variants.

The existence of acapsular *S.* Typhi variants in human infection sites had been reported before this study. Acapsular *S.* Typhi variants lack the entire *viaB* locus or genes in the locus ^50,51^. For instance, in Pakistan, 13 of the 102 clinical isolates ^50^ and 2 of the 35 clinical isolates ^51^ were found to be acapsular variants. Nothing was known regarding the emergence of other capsule variants. Here we discovered that three new pathogenic variants of capsulated bacteria, the so-called “hypo-capsule”, “super-capsule”, and “hyper-capsule” variants, have different strengths in their pathogenicity. The “hypo-capsule” *S.* Typhi Vi variants have a markedly increased infectivity whose serum survival was almost comparable to WT *S.* Typhi. The “hyper-capsule” *S.* Typhi variants are superb in evading the host innate immune system and colonizing the gallbladder and other organs, as a 100,000-fold increased bacterial burden was observed. Both the bacteria-associated hyper capsule and the shed Vi free from the bacteria are vital for the enhanced virulence of the hyper-capsule variants. It is also notable that the effects associated with the shed Vi is enhancing co-infected pathogens. The “super-capsule” *S.* Typhi variants diverge into two subgroups: some variants phenocopy WT and the other phenocopy hyper-capsule variants.

The most striking finding of this study is that many of the frequent clinical missense mutations in *tviD* and *tviE* are hyper-capsule variants. Hyper-capsule *S*. Typhi strains were resistant to human neutrophil-mediated phagocytosis. In mice, hyper-capsule *S*. Typhi induced liver abscesses, clinical signs (overall weakness, eye discharge, and diarrhea), and deaths in 50% of infected mice. Intriguingly, all mice infected with hyper-capsule variants had *S.* Typhi colonized in their gallbladder. Recent genome sequencing analyses of *S.* Typhi gallbladder isolates reported 3 of the hyper-capsule missense mutations that we found in this study: *tviD* R508H, *tviE* C137R, and *tviE* A462V ^33,65^, suggesting a possible correlation between hyper-capsule variants and heightened gallbladder colonization in humans.

The link between hyper-capsule and hypervirulence was documented in other capsulated pathogens, including *Klebsiella pneumonia*, *Acinetobacter baumanii,* and *Streptococcus pneumoniae* ^6,67–71^, which seems to point to parallel evolution in capsulated pathogens. The molecular bases for their hyper-capsule phenotypes appear to be more complex than that of *S.* Typhi, as deletions, insertions, and several non-synonymous SNPs were involved ^6,72–74^. The most similar adaptation evolution to that of *S.* Typhi is *K. pneumoniae*, as SNPs in the capsule biosynthesis gene *wzc* lead to hyper-capsule production, which confers phagocytosis resistance, enhanced dissemination, and increased mortality in animal models ^6,67^. In human infection sites, diarrhea and overall weakness are known general typhoid fever clinical signs. Although whether hyper-capsule *S*. Typhi was involved is unknown, typhoid fever cases with liver abscess, in some cases, with gallbladder colonization, have been reported ^75–80^.

We have made several technical advancements through the development and refinement of novel molecular tools and methodologies. We have effectively created an *in vivo* molecular tool that enables us to examine the impact of clinical missense mutations on the mode of action for TviE. TviE studies, as well as our research utilized TviD C127R and two experimental mutations, illustrate the distinct correlation between amino acid substitutions found in clinical missense mutations and the hypo-capsule or hyper-capsule phenotypes observed in TviE and TviD variants. We note that the mechanistic studies conducted by TviD are comparatively less comprehensive in comparison to those conducted by TviE. In contrast to TviE mutations, the mutations in TviD were observed to occur throughout its 3D structure, which presents challenges in identifying a discernible adaptive evolution theme. However, the complexity of this situation is consistent with the prediction that mutations affect various stages of Vi production. We predict that these stages include: (a) mutations that directly impact the catalytic activity, either inhibiting or promoting the reaction (e.g., C127R); (b) mutations that affect catalysis during the pre- or post-stage, involving the reception of the glycan substrate or the transfer of the resulting glycan to the next enzyme, often found in the loop region; and (c) mutations that influence the formation of multimers or multienzyme complexes, typically located on the surface. Further investigations are warranted to fully understand these effects by specific clinical mutations. We have developed a whole-genome sequencing (WGS) analysis pipeline for *S.* Typhi. It is expected that the murine model used, combined with hyper-capsule *S.* Typhi, will prove to be a valuable *S.* Typhi infection model for the initial phase of infection. This model will facilitate the detection of variations in virulence during the early days of infection. However, it is crucial to recognize the necessity for additional research utilizing more appropriate animal models in order to investigate the effects of capsule variants during the later stages of infection.

In summary, by combining epidemiology, genomics, and molecular investigation, here we provide much-needed valuable insights into the various types and consequences of the Vi capsule variants, as well as a future outlook on the occurrence and dissemination of the capsule variants of *S.* Typhi.

## Online Methods

### Bacterial strains

*S*. Typhi ISP2825 ^52^ was used as a parent strain for generating the following strains: *ΔtviBC*, *ΔtviD,* Δ*tviE-vexE* (a deletion of *tviE, vexA, vexB, vexC, vexD,* and *vexE*)*, tviD* A35S, *tviD* C127R, *tviD* E236K, *tviD* H253A, *tviD* T293A, *tviD* T424I, *tviD* R508H, *tviD* R523H, *tviD* G616S, *tviD* S676N, *tviD* G752C, *tviD* D763N, *tviD* E810K, *tviE* C137R, *tviE* Q219R, *tviE* P263S, *tviE* K266N, *tviE* K266R, *tviE* K266E, *tviE* S290G, *tviE* S290R, *tviE* A421T, *tviE* A462V, *tviE* V495A, *tviA* V62I, *tviD* C127S, *tviD* C127A, WT (Cm^R^), *tviE* P263S (Kan^R^), and *tviE* K266N (Kan^R^) (Supplementary Table 7). We note that we used gene names (e.g., *tviD*) with the protein code (e.g., C to R) and protein residue location (e.g., 127) to reflect missense mutations (and distinguish them from synonymous single-point mutations [SNPs]). Genetic manipulations were performed as described previously ^59,81^. In brief, pSB890 (Tet^R^ with *sacB* for sucrose selection), a suicide vector, was used as a backbone to introduce point mutations and epitope tags. The vector pSB890 was digested with restriction enzymes (BamHI-HF [NEB, cat# R3136S] and NotI-HF [NEB, cat# R3189S]). Inserts for each mutant were amplified by conducting PCR reactions using Herculase® II Fusion DNA Polymerase (Invitrogen, cat# 600679) and specific primers (Supplementary Table 7) with the *S*. Typhi ISP2825 genome as a template. The digested vector and inserts were Gibson-assembled (T5 exonuclease, NEB cat# M0363S; Phusion polymerase, NEB cat# M0530S; Taq DNA ligase, NEB cat# M0208L) to generate the required plasmids (Supplementary Table 7). The resulting plasmids were transformed into *E. coli* ß2163 Δnic35 for conjugation and subsequent homologous recombination in *S.* Typhi. All strains were verified by performing Sanger sequencing available through the Cornell Institute Biotechnology Resource Center (BRC) Genomics Facility (RRID: SCR_021727) or Eton Bioscience (RRID: SCR_003533).

### Mammalian cell culture conditions

Human epithelial cell Henle-407 was cultured in DMEM high glucose (Invitrogen) supplemented with 10% FBS (Hyclone cat# SH30396.03, Lot# AD14962284). Sialic acid contents of the FBS used were validated, which was ∼99% Neu5Ac and less than 1% Neu5Gc. Cells were kept at 37°C in a cell culture incubator with 5% CO_2_. Mycoplasma testing was conducted regularly as part of the cell maintenance practice.

### *S*. Typhi whole genome sequence analyses

A bioinformatics pipeline was established for WGS comparison analyses of 5,379 *S*. Typhi clinical isolates. In brief, all *S*. Typhi WGSs that were available in the NCBI’s GenBank database (5,749 strains as of 11/03/21) were re-annotated using the Prokaryotic Genome Annotation Pipeline (PGAP, 2021-07-01.build5508; RRID: SCR_021329) to synchronize all the annotations under same criteria. RefSeq’s gene numbers were used to identify the gene pool from each strain and a single digit code was allocated to each gene: -1, absence of gene; 0, predicted pseudogene; 1, presence of gene, using a bash script. Digit code was left blank if the gene was partially sequenced (Supplementary Table 1). To ensure that the dataset only includes *S.* Typhi WGSs, the following two additional filters were used: 1) SeqSero2 ^46^ and 2) typhoid toxin subunits (*cdtB*, *pltA*, and *pltB*) ^47^, resulting in a total of 5,379 *S.* Typhi strains. Point mutations on each Vi synthesis gene were examined by using Clustal Omega (RRID: SCR_001591) multiple sequence alignments and visualized with Prism 9.4.1 (GraphPad, RRID: SCR_002798). WGS accession numbers and the digit codes of target genes used in this study are described in Supplementary Table 1.

For epidemiological studies, *S*. Typhi WGSs from the NCBI’s GenBank database (5,749 strains as of 11/03/21) and the Pathogenwatch (13,081 strains as of 12/27/23) were combined, and duplicates were removed. The *S*. Typhi WGSs of 16,368 strains were analyzed by genotyphi (https://github.com/typhoidgenomics/genotyphi) to define their lineages and antimicrobial resistance of ciprofloxacin and azithromycin. *tviE* and *tviD* nucleotide sequences were extracted using SeqKit ^82^, and single nucleotide polymorphisms (SNPs) of *tviE* and *tviD* were identified with a custom bash script. The data were visualized with Prism 9.4.1 (GraphPad, RRID: SCR_002798).

### Immunoblots

*S*. Typhi ISP2825 was grown in Luria-Bertani (LB) broth at 37 °C overnight. Fifty microliters of the culture were inoculated in 2 ml of 86 mM NaCl LB, and cultured until OD_600_ reaches 0.9-1.0. A volume equivalent to OD_600_ = 1.0 was pelleted at 5,000 ×g for 5 min and washed with PBS. When the shed Vi was indicated, 0.5 ml of the culture supernatant was saved after the aforementioned pelleting, which was further centrifuged at 20,000 ×g for 10 min. Two hundred microliters of the supernatant were collected; 40 µl of 6x SDS sample buffer (360 mM Tris-HCl, 12% SDS, 60% Glycerol, 5% ß-mercaptoethanol, and 0.2% Bromophenol blue) containing 20 µg/ml proteinase K was added. Note that we used proteinase K for glycan immunoblots. When the Vi expression of *S*. Typhi was compared to the shed Vi in the supernatant, the bacterial pellet was resuspended in 1 mL of 1x SDS sample buffer containing 20 µg/mL proteinase K, followed by vortexing at 15 minutes intervals for 1 hour at 37°C. In most experiments that did not need to quantify the amounts of the shed Vi, the bacterial pellet was resuspended in 0.4 mL of 1x SDS sample buffer containing 20 µg/mL proteinase K. For protein immunoblots (Myc and Flag), the pellets were lysed in 0.4 mL of 1x SDS sample buffer without proteinase K, followed by boiling at 98°C for 5 min. The lysates were centrifuged at 20,000 ×g for 10 minutes; 5 µL (1 x 10^7^ cells when 0.4 ml of 1x SDS buffer was used) of the lysates was run on 8–16% Mini-PROTEAN® TGX™ Precast Protein Gels (Bio-Rad, cat# 4561106) for Vi and O9 detection or 8% Tris-Glycine SDS-PAGE gel for V5, Strep, Myc, Flag, and HA detection. The SDS-PAGE gels were wet-transferred at 20 V overnight at 4 °C using Bjerrum Schafer-Nielsen transfer buffer (48 mM Tris, 39 mM glycine, and 20% methanol) to nylon membranes (Biodyne B Nylon Membrane, Pall) for Vi and LPS or nitrocellulose membrane (Amersham, cat# 10600002)) for protein samples. *Salmonella* Vi antiserum (pan-Vi antibody) at 1/300 dilution (BD Difco, cat# 228271, lot# 1357166, RRID: AB_2934033), anti-Vi monoclonal antibody (*O*Ac-Vi antibody) at 1/200 dilution (SSIDiagnostica, cat# REF15514, lot# 188O-3, RRID: AB_2934031), or rabbit anti-O9 antiserum at 1/200 dilution (SSIDiagnostica, cat# REF40276, lot# 636O-H5, RRID: AB_2934032) was used to detect Vi or O9 antigens. SuperSignal™ West Femto Maximum Sensitivity Substrate (ThermoFisher, cat# 34095) was used to develop the images using iBright™ CL1500 Imaging System (ThermoFisher). Vi and O9 intensities were calculated using iBright™ analysis software (ThermoFisher, RRID: SCR_017632), and quantified using Prism 9.4.1 (GraphPad, RRID: SCR_002798).

### TviE and TviD structure predictions

TviE or TviD structure was predicted using RoseTTAFold ^55^ and AlphaFold v2 ^54^ or AlphaFold v2, respectively. Mapping the clinical missense point mutations, electrostatic potential calculation, and superimposition of the predicted TviE and TviD structures from AlphaFold v2 were performed using ChimeraX (University of California at San Francisco, RRID: SCR_01587).

### TviE-inducible system

#### TviE-inducible system construction

*S*. Typhi ISP2825 lacking *tviE-vexA-vexB-vexC-vexD-vexE* (Δ*tviE-vexE*) was constructed and transformed with pBAD-*tviE*-Flag carrying *tviE* sequence of WT, P263S (hyper Vi), K266N (hypo Vi), or S290R (hyper Vi) (Supplementary Table 7).

#### TviE mutant characterization

*S*. Typhi ISP2825 Δ*tviE-vexE* (pBAD-*tviE*-Flag) was cultured overnight in LB containing 100 µg/mL ampicillin and 0.5% glucose. Two hundred fifty microliters of the overnight culture were transferred into 10 mL of LB containing 86 mM NaCl, 100 µg/mL ampicillin, and 0.5% glucose and cultured until OD_600nm_ reaches 0.9-1.0. A control sample (at 0 min, without L-arabinose induction) was collected at this step (a volume equivalent to OD_600nm_ = 1). The culture was centrifuged at 5,000 ×g at 4°C for 10 minutes and resuspended in 9 mL of LB containing 86 mM NaCl, 100 µg/mL ampicillin, and 0.1% L-arabinose; the culture was aliquoted into four tubes (2 mL each) and cultured. A volume equivalent to OD_600nm_ = 1 was collected and pelleted at each indicated time point (15, 30, 45, and 60 minutes). Immunoblots were performed as described above.

### Bacterial growth/overall fitness determinations

The overnight culture was 10-fold serial-diluted, and further diluted in PBS to prepare 8 × 10^6^ cells/ml (corresponding to OD_600nm_ = 0.01). Ten microliters of the prepared dilution (8 x 10^4^ cells) were added to 150 µL of 86 mM NaCl LB or 300 mM NaCl LB on a 96-well plate; OD_600nm_ was measured at 37°C at 30 min intervals for 24 hours; the plate was shaken for 5 seconds before each measurement.

### Image cytometry for Vi and O9 expressions

*S*. Typhi ISP2825 and the indicated each single amino acid point mutant was grown in LB overnight; 50 µL of the overnight culture was inoculated into 2 mL of 86 mM NaCl LB and cultured until OD_600nm_ reaches 0.9-1.0. A volume equivalent to OD_600nm_ = 1.0 was pelleted at 5,000 ×g for 5 minutes and resuspended in PBS. The suspension was pelleted at 5,000 xg for 5 minutes, re-suspended in PBS containing 4% paraformaldehyde (PFA), and incubated for 1 hour. The lack of viability of *S*. Typhi was confirmed by spreading and culturing the fixed cells on LB plates. In parallel, during the fixation step, coverslips were coated with poly-D-lysine (Gibco, cat# A3890401) for 1 hour, washed with Milli-Q water, and dried. In one hour, the fixed cells were pelleted at 5,000 ×g for 5 minutes and re-suspended in 100 µL of PBS. The samples were placed on a poly-D-lysine-coated coverslip for 30 minutes; the coverslip was washed with PBS, incubated in PBS containing 3% BSA for 30 minutes for blocking, replaced the blocking buffer with PBS/3% BSA containing *O*Ac-Vi antibody (1:200) and O9 antibody (1:200), and incubated for 1 hour. In one hour, the coverslip was washed with PBS, replaced the buffer with PBS/3% BSA containing Alexa-488 goat anti-rabbit (Invitrogen, cat# A11034, RRID: AB_2576217, 1:4000) and Alexa-594 goat anti-mouse antibody (Invitrogen, cat# A11032, RRID: AB_2534091, 1:4000), and incubated for 30 minutes. Using a BZ-X810 (Keyence) microscope, *S*. Typhi was imaged under Plan Apochromat 60X Oil objective (BZ-PA60); we set the Keyence imaging program to automatically acquire and stitch 49 images (7 by 7), resulting in one overall image covering the 1.256 mm × 0.942 mm area of each coverslip. Alexa-Flour 594 brightness (Vi) of each of the acquired overall images was quantified; we let the BZ-X810 image analyzer find the area that had *S.* Typhi O9 staining and quantified the brightness for Vi. We note that the +5 expand option in BZ-X810 Analyzer was used given the location of Vi and LPS O9 on the surface of *S.* Typhi. The total Alexa-Flour 594 brightness (Vi) was then divided by the number of *S*. Typhi counted from O9 staining to determine the average Vi brightness/cell from the stitched image. All image cytometry processes of these fixed cells were performed at room temperature.

### Alcian blue staining

*S*. Typhi WT, Δ*tviBC*, *tviD* C127R, and *tviE* K266N were grown in LB broth at 37°C overnight. Fifty microliters of the culture were inoculated in 2 mL of 86 mM NaCl LB, and cultured until OD_600_ reaches 0.9-1.0. A volume equivalent to OD_600_ = 1.0 was pelleted at 5,000 ×g for 5 minutes and washed with PBS. The pellet was incubated with 200 µL of alcian blue staining solution (1% Alcian blue, and 3% acetate) for 15 minutes, washed with 3% acetate four times, lysed with 250 µL of lysis solution (1% SDS, 100 mM Tris pH 7.5), and heated at 95°C for 5 minutes. After centrifugation at 20,000 ×g for 10 minutes, 200 µL of the supernatant was transferred to a 96-well plate. The absorbance at 620 nm was measured using a Tecan Infinite 200 Pro microplate reader (Tecan).

### Invasion and adhesion assays

Henle-407 cells (5 x 10^4^) were seeded on a 24-well plate a day before infection. For imaging intracellular bacteria, a coverslip was placed on 24 well plates before seeding the cells. The cells were infected with *S.* Typhi WT or mutant in Hanks’ Balanced Salt Solution (Invitrogen) at 15 multiplicity of infection (m.o.i.) for 1 hour, washed with PBS, and incubated with 100 µg/mL gentamicin for 45 min to kill extracellular bacteria.

For colony forming unit (CFU) determination at 2 hours post-infection, the infected cells were washed with PBS, lysed in 1 mL of PBS containing 0.1% sodium deoxycholate for 15 minutes at room temperature, and 100 µL of 10^-1^-diluted lysate was spread on LB agar plates. The cells for the 24 hours post-infection were maintained in complete DMEM containing 10 µg/mL gentamicin. Colonies were counted to calculate the total number of intracellular bacteria.

For immunofluorescence assays, the coverslips were fixed in PBS/4% PFA overnight at 4°C, washed with Tris-buffered saline (TBS) to quench PFA, and permeabilized for 30 minutes with PBS containing 3% BSA, 0.2% Triton X-100, and 10 mM Tris. For imaging intracellular *S*. Typhi after 2 hours post-infection, the coverslips were incubated with anti-*Salmonella* antibody (Difco, cat# 240993, 1:4000) and Alexa-Fluor 488 Phalloidin (Invitrogen, cat# A12379) for 1 hour, washed with PBS-T (PBS/0.1% Tween 20), incubated with Alexa Fluor-594-labeled anti-rabbit antibody for 30 minutes, and counterstained the nuclei with 4’,6-diamidino-2-phenylindole (DAPI, Invitrogen, cat# D3571) for 10 minutes. Intracellular *S*. Typhi was imaged using a BZ-X810 microscope (Keyence); Plan Fluorite 20X LD PH (BZ-PF20LP) objective was used to acquire 100 images covering 5.290 mm × 3.968 mm area of each coverslip. We let the BZ-X810 image analyzer find host cells (Phalloidin+) and count intracellular *S*. Typhi; the total counted number of *S*. Typhi was divided by the number of DAPI-stained nuclei to determine *S*. Typhi/cell from each image. The fully-focused images presented in Fig. 3j were acquired using Plan Apochromat 60X Oil (BZ-PA60) objective.

For inside-outside staining presented in Fig. 3l-m (1-hour post-infection), the fixed cells on coverslips were incubated with PBS/3% BSA for 30 minutes before incubating with anti-*Salmonella* antibody for 2 hours to stain extracellular, attached bacteria. The coverslips were washed with PBS and incubated with Alexa Fluor-488-labeled anti-rabbit antibody for 1 hour. The coverslips were then washed with PBS and permeabilized for 30 minutes with PBS containing 3% BSA, 0.2% Triton X-100, and 10 mM Tris. To stain both intracellular and extracellular, attached bacteria, the coverslips were incubated with anti-*Salmonella* antibody for 1 hour, washed with PBS-T, incubated with Alexa Fluor-594-labeled anti-rabbit antibody for 1 hour, and the nuclei were counterstained with DAPI. All the processes were performed at room temperature except PFA fixation.

### Image acquisition and quantification

Immunofluorescent and agglutination images were acquired by using a BZ-X810 microscope (Keyence). The filters used in this study are as follows: Alexa-Fluor 488 and Alexa-Fluor 488 Phalloidin, 470/40 nm excitation with 525/50 nm emission (OP-87763); Alexa-Fluor 594, 560/40 nm excitation with 630/75 nm emission (OP-87765); DAPI, 360/40 nm excitation with 460/50 nm emission (OP-87762). The immunofluorescent images of intracellular *S*. Typhi were quantified with the ‘Hybrid Cell Count’ function of the BZ-X810 image analyzer. For quantifying *S.* Typhi aggregates in agglutination assays, CellProfiler Image Analysis Software (BROAD institute, RRID: SCR_007358) was utilized. For Vi quantification, the membranes were imaged using an iBright CL1500 imager (ThermoFisher) and analyzed with iBright analysis software 5.1.0 (ThermoFisher, RRID: SCR_017632). The length differences of Vi were analyzed using ImageJ 1.53k (National Institutes of Health, RRID: SCR_003070). All statistics were performed using Prism 9.4.1 (GraphPad, RRID: SCR_002798).

### Serum resistance assay

#### Human sera

The process of collecting human blood samples was carried out in accordance with the protocols that have been approved by the Institutional Review Board for Human Participants at Cornell University. A nurse practitioner at the Cornell Human Metabolic Research Unit conducted a peripheral blood draw to obtain primary neutrophils and sera. The data were analyzed in an anonymous manner. All adult participants provided informed consent. The written consent form was provided to all participants. De-identified human blood samples were clotted for 1 hour at room temperature, and centrifuged at 2,000 ×g. The supernatant (healthy human serum) was harvested, aliquoted, and stored at -80°C until use. When indicated, a serum aliquot was incubated at 56°C for 1 hour to prepare complement-inactivated human serum (iHS).

#### Serum resistance assay

Overnight *S*. Typhi culture was prepared as described above; 50 µL of the culture was inoculated into 2 mL of 86 mM NaCl LB and incubated at 37°C until OD_600nm_ reaches 0.9-1.0. A volume equivalent to OD_600nm_=1.0 was centrifuged at 5,000 ×g for 5 minutes. The pellet was resuspended in 0.5 mL PBS, after which 10 µl of *S*. Typhi (1.6 x 10^7^ cells) was added into 150 µL PBS (1.6 x 10^7^ cells/ 160 µL). For serum resistance assays, 10 µL of *S*. Typhi in PBS (1 x 10^6^ cells) was added into 90 µL of PBS in the absence and presence of human serum or inactivated human serum. The samples were incubated for 2 hours at 37°C. CFU was determined by culturing 10-fold serial dilutions on LB agar plates; colonies were counted after 16 hrs. Plate images were acquired using an iBright™ CL1500 Imaging System (ThermoFisher).

### Vi purification

*S*. Typhi WT, Δ*tviBC*, or *tviE* P263S was grown in LB broth at 37°C overnight. Two hundred fifty microliters of the culture were inoculated in 10 mL of 86 mM NaCl LB, and cultured until OD_600_ reaches 1.5. The culture was centrifuged at 4,000 rpm for 20 minutes at 4°C. The supernatant of each strain was 0.22 µm-filtered, and incubated with DNase I and Rnase A for 2 hours at 37°C, followed by incubating with Proteinase K for 2 hours at 37°C. The supernatant (10 mL) was 0.22 µm-filtered before being concentrated to 0.5 mL using a 30 kDa cut-off filter. The resulting 0.5 mL was mixed with 10 mL PBS before being passed through the 30 kDa cut-off filter to bring the volume to 0.5 mL. This washing-concentration procedure was repeated 6 times to ensure that the enzymes and other contaminants were removed. The resulting samples (100 µL) were incubated with anti-O9-conjugated agarose beads for 1 hour at RT to minimize LPS contamination. The Vi was further-purified with size exclusion chromatography (AKTA) using Superdex 200 Increase 10/300 GL column (Cytiva, cat# 28-9909-44), and quantified by conducting immunoblots.

### Animal experiments

Mice experiments were conducted according to a protocol approved by Cornell IACUC. Sex-matched groups of 7-to 11-week-old Cmah-null mice expressing the human-type glycan receptor (Jackson Laboratory, RRID: IMSR_JAX:017588) ^47,57,59,60,63,83–85^ were infected intraperitoneally with 8 x 10^5^ *S*. Typhi WT, *tviE* K266N, or *tviE* P263S strain in 100 µL of PBS. Infected mice were sacrificed, and organs (liver, spleen, and gallbladder) were harvested 3 days after infection, except two mice in *tviE* P263S group that died on Day 1. The organs were mechanically homogenized with PBS/0.05% sodium deoxycholate; 5 mL, 3 mL, or 2 mL of lysis buffer was used for the liver, spleen, and gallbladder, respectively. CFUs were determined by plating 10-fold serial dilutions of homogenates on LB agar plates. The number of total CFU in each organ was calculated by multiplying dilution factors.

For co-infection experiments of WT with *tviE* P263S or *tviE* K266N, 4 x 10^5^ WT *S*. Typhi (Chloramphenicol-resistant) and 4 x 10^5^ *S*. Typhi *tviE* P263S (Kanamycin-resistant) or K266N (Kanamycin-resistant) were prepared in 100 µL of PBS (8 x 10^5^ in total) for infection. Organ CFUs were assessed by plating the diluted lysates on LB agar plates containing 2.5 µg/mL Chloramphenicol for WT or 50 µg/mL Kanamycin for *tviE* P263S and K266N. For WT *S.* Typhi infection experiments with shed Vi, the amount of shed Vi corresponding to the surface Vi of ∼1.6 x 10^9^ *S*. Typhi was mixed with 8 x 10^5^ *S*. Typhi WT in 100 µL of PBS for infection. *S.* Typhi Δ*tviBC* preparations having comparable LPS concentrations were used as a control.

### Measurement of ROS burst from human peripheral blood neutrophils

Human peripheral blood neutrophils from human blood samples were isolated as described elsewhere. Briefly, 10 mL of peripheral blood were collected in a EDTA-treated tube, and gently mixed with 5 mL of 3% gelatin (Difco, cat# 214340)/0.1% glucose/0.9% NaCl. The red blood cells were sedimented at RT for 45 minutes. The neutrophil-rich supernatant was transferred onto the top of 3 mL of Ficoll-Paque Premium (GE Healthcare, cat# 17-5442-02) in a 15 mL conical tube. The samples were centrifuged at 1,500 rpm for 20 min at 16°C. The pellet was resuspended in 6 mL of red blood cell lysis buffer (1 mL of PBS and 5 mL of distilled water) and incubated for 45 seconds, followed by adding 2 mL of 3% NaCl solution to stop the reaction. The neutrophil samples were centrifuged at 1,500 rpm for 5 minutes at 4°C, and resuspended in 1 mL of 2% FBS/phenol red-free RPMI-1640. *S*. Typhi WT and capsule variants were grown in 2 mL of 86 mM NaCl LB until OD_600_ reaches 1.0. Each strain was opsonized with human sera in PBS for 30 minutes at RT. 5 x 10^4^ neutrophils in 90 µL of phenol-free RPMI (ThermoFisher, cat# 11835030) supplemented 2% FBS with 1 mM luminol (Sigma, cat# 123072) were seeded into a black opaque 96-well plate, and 10 µL of opsonized *S*. Typhi (5 x 10^5^/10 µL) were added (10 m.o.i.). Luminescence was recorded at 2-minute interval using Tecan Infinite 200 Pro microplate reader (Tecan).

### Quantification and statistical analysis

Data were tested for statistical significance with the GraphPad Prism software. The number of replicates for each experiment and the statistical test performed are indicated in the figure legends. Image analysis and quantification were performed using ImageJ. The number of biological replicates and the statistical method are described in each figure legend. At least 3 independent experiments were performed throughout the study.

## Author contributions

G. Y. L. conceptualized this research, executed all experiments, interpreted the results, and wrote the manuscript. J.S. conceptualized this research, acquired the funding, interpreted the results, executed animal experiments, and wrote the manuscript.

## Supplementary information

Supplemental information (Supplementary Figs. 1-3 and Tables 1-7) can be found online.

**Correspondence and requests for materials** should be addressed to Jeongmin Song.

## Materials availability

*Salmonella* mutant strains used for this study would be made available to other researchers via the Institutional Material Transfer Agreements.

## Data and code availability

The published article includes all datasets generated during this study.

## REFERENCES

1 Follador, R. et al. The diversity of Klebsiella pneumoniae surface polysaccharides. Microb Genom 2, e000073, doi:10.1099/mgen.0.000073 (2016).

2 Lee, S. et al. Glycan-mediated molecular interactions in bacterial pathogenesis. Trends Microbiol 30, 254–267, doi:10.1016/j.tim.2021.06.011 (2022).

3 Mostowy, R. J. & Holt, K. E. Diversity-Generating Machines: Genetics of Bacterial Sugar-Coating. Trends Microbiol 26, 1008–1021, doi:10.1016/j.tim.2018.06.006 (2018).

4 O’Riordan, K. & Lee, J. C. Staphylococcus aureus capsular polysaccharides. Clin Microbiol Rev 17, 218–234, doi:10.1128/cmr.17.1.218-234.2004 (2004).

5 Paton, J. C. & Trappetti, C. Streptococcus pneumoniae Capsular Polysaccharide. Microbiol Spectr 7, doi:10.1128/microbiolspec.GPP3-0019-2018 (2019).

6 Ernst, C. M. et al. Adaptive evolution of virulence and persistence in carbapenem-resistant Klebsiella pneumoniae. Nat Med 26, 705–711, doi:10.1038/s41591-020-0825-4 (2020).

7 Sande, C. & Whitfield, C. Capsules and Extracellular Polysaccharides in Escherichia coli and Salmonella. EcoSal Plus 9, eESP00332020, doi:10.1128/ecosalplus.ESP-0033-2020 (2021).

8 Wetter, M. et al. Molecular characterization of the viaB locus encoding the biosynthetic machinery for Vi capsule formation in Salmonella Typhi. PLoS One 7, e45609, doi:10.1371/journal.pone.0045609 (2012).

9 WHO. Typhoid vaccines: WHO position paper, March 2018 - Recommendations. Vaccine, doi:10.1016/j.vaccine.2018.04.022 (2018).

10 Lesinski, G. B. & Westerink, M. A. Vaccines against polysaccharide antigens. Curr Drug Targets Infect Disord 1, 325–334, doi:10.2174/1568005014605964 (2001).

11 Dahora, L. C. et al. Salmonella Typhi Vi capsule prime-boost vaccination induces convergent and functional antibody responses. Sci Immunol 6, eabj1181, doi:10.1126/sciimmunol.abj1181 (2021).

12 Hashimoto, Y., Li, N., Yokoyama, H. & Ezaki, T. Complete nucleotide sequence and molecular characterization of ViaB region encoding Vi antigen in Salmonella typhi. J Bacteriol 175, 4456–4465, doi:10.1128/jb.175.14.4456-4465.1993 (1993).

13 Virlogeux, I., Waxin, H., Ecobichon, C. & Popoff, M. Y. Role of the viaB locus in synthesis, transport and expression of Salmonella typhi Vi antigen. Microbiology (Reading) 141 **( Pt** **12**), 3039–3047, doi:10.1099/13500872-141-12-3039 (1995).

14 Seth-Smith, H. M. SPI-7: Salmonella’s Vi-encoding Pathogenicity Island. J Infect Dev Ctries 2, 267–271, doi:10.3855/jidc.220 (2008).

15 Pickard, D. et al. Characterization of defined ompR mutants of Salmonella typhi: ompR is involved in the regulation of Vi polysaccharide expression. Infect Immun 62, 3984–3993, doi:10.1128/iai.62.9.3984-3993.1994 (1994).

16 Winter, S. E. et al. The TviA auxiliary protein renders the Salmonella enterica serotype Typhi RcsB regulon responsive to changes in osmolarity. Mol Microbiol 74, 175–193, doi:10.1111/j.1365-2958.2009.06859.x (2009).

17 Virlogeux, I., Waxin, H., Ecobichon, C., Lee, J. O. & Popoff, M. Y. Characterization of the rcsA and rcsB genes from Salmonella typhi: rcsB through tviA is involved in regulation of Vi antigen synthesis. J Bacteriol 178, 1691–1698, doi:10.1128/jb.178.6.1691-1698.1996 (1996).

18 Zhang, H., Zhou, Y., Bao, H. & Liu, H. W. Vi antigen biosynthesis in Salmonella typhi: characterization of UDP-N-acetylglucosamine C-6 dehydrogenase (TviB) and UDP-N-acetylglucosaminuronic acid C-4 epimerase (TviC). Biochemistry 45, 8163–8173 (2006).

19 Wear, S. S., Sande, C., Ovchinnikova, O. G., Preston, A. & Whitfield, C. Investigation of core machinery for biosynthesis of Vi antigen capsular polysaccharides in Gram-negative bacteria. J Biol Chem 298, 101486, doi:10.1016/j.jbc.2021.101486 (2022).

20 Liston, S. D., Ovchinnikova, O. G. & Whitfield, C. Unique lipid anchor attaches Vi antigen capsule to the surface of Salmonella enterica serovar Typhi. Proc Natl Acad Sci U S A 113, 6719–6724, doi:10.1073/pnas.1524665113 (2016).

21 Crump, J. A., Luby, S. P. & Mintz, E. D. The global burden of typhoid fever. Bulletin of the World Health Organization 82, 346–353 (2004).

22 Crump, J. A. & Mintz, E. D. Global trends in typhoid and paratyphoid Fever. Clin Infect Dis 50, 241–246, doi:10.1086/649541 (2010).

23 Parry, C. M., Hien, T. T., Dougan, G., White, N. J. & Farrar, J. J. Typhoid fever. N Engl J Med. 347, 1770–1782. (2002).

24 Parry, C. M. & Threlfall, E. J. Antimicrobial resistance in typhoidal and nontyphoidal salmonellae. Curr Opin Infect Dis 21, 531–538, doi:10.1097/QCO.0b013e32830f453a (2008).

25 Everest, P., Wain, J., Roberts, M., Rook, G. & Dougan, G. The molecular mechanisms of severe typhoid fever. Trends Microbiol 9, 316–320 (2001).

26 Monack, D. M. Helicobacter and salmonella persistent infection strategies. Cold Spring Harb Perspect Med 3, a010348, doi:10.1101/cshperspect.a010348 (2013).

27 Lovane, L. et al. Carriage prevalence of Salmonella enterica serotype Typhi in gallbladders of adult autopsy cases from Mozambique. J Infect Dev Ctries 10, 410–412 (2016).

28 Gunn, J. S. et al. Salmonella chronic carriage: epidemiology, diagnosis, and gallbladder persistence. Trends Microbiol 22, 648–655 (2014).

29 Monack, D. M. Salmonella persistence and transmission strategies. Curr Opin Microbiol 15, 100–107, doi:10.1016/j.mib.2011.10.013 (2012).

30 Blaser, M. J. & Kirschner, D. The equilibria that allow bacterial persistence in human hosts. Nature 449, 843–849, doi:10.1038/nature06198 (2007).

31 Gibbs, K. D. et al. Human variation impacting MCOLN2 restricts Salmonella Typhi replication by magnesium deprivation. Cell Genom 3, 100290, doi:10.1016/j.xgen.2023.100290 (2023).

32 Crawford, R. W. et al. Gallstones play a significant role in Salmonella spp. gallbladder colonization and carriage. Proc Natl Acad Sci U S A 107, 4353–4358, doi:10.1073/pnas.1000862107 (2010).

33 Gonzalez-Escobedo, G., Marshall, J. M. & Gunn, J. S. Chronic and acute infection of the gall bladder by Salmonella Typhi: understanding the carrier state. Nat Rev Microbiol 9, 9–14, doi:10.1038/nrmicro2490 (2011).

34 Song, J., Gao, X. & Galan, J. E. Structure and function of the Salmonella Typhi chimaeric A(2)B(5) typhoid toxin. Nature 499, 350–354, doi:10.1038/nature12377 (2013).

35 Song, J. et al. A mouse model for the human pathogen Salmonella typhi. Cell Host Microbe 8, 369–376, doi:10.1016/j.chom.2010.09.003 (2010).

36 Wangdi, T. et al. The Vi capsular polysaccharide enables Salmonella enterica serovar typhi to evade microbe-guided neutrophil chemotaxis. PLoS Pathog 10, e1004306, doi:10.1371/journal.ppat.1004306 (2014).

37 Galán, J. E. Common themes in the design and function of bacterial effectors. Cell Host Microbe 5, 571–579 (2009).

38 Nikolaus, T. et al. SseBCD proteins are secreted by the type III secretion system of Salmonella pathogenicity island 2 and function as a translocon. J Bacteriol 183, 6036–6045 (2001).

39 Srikanth, C. V., Mercado-Lubo, R., Hallstrom, K. & McCormick, B. A. Salmonella effector proteins and host-cell responses. Cell Mol Life Sci 68, 3687–3697 (2011).

40 Abrahams, G. L. & Hensel, M. Manipulating cellular transport and immune responses: dynamic interactions between intracellular Salmonella enterica and its host cells. Cell Microbiol 8, 728–737, doi:10.1111/j.1462-5822.2006.00706.x (2006).

41 Elhadad, D., McClelland, M., Rahav, G. & Gal-Mor, O. Feverlike Temperature is a Virulence Regulatory Cue Controlling the Motility and Host Cell Entry of Typhoidal Salmonella. J Infect Dis 212, 147–156 (2015).

42 Wilson, R. P. et al. The Vi capsular polysaccharide prevents complement receptor 3-mediated clearance of Salmonella enterica serotype Typhi. Infect Immun 79, 830–837, doi:10.1128/iai.00961-10 (2011).

43 Hiyoshi, H. et al. Mechanisms to Evade the Phagocyte Respiratory Burst Arose by Convergent Evolution in Typhoidal Salmonella Serovars. Cell Rep 22, 1787–1797, doi:10.1016/j.celrep.2018.01.016 (2018).

44 Hart, P. J. et al. Differential Killing of Salmonella enterica Serovar Typhi by Antibodies Targeting Vi and Lipopolysaccharide O:9 Antigen. PLoS One 11, e0145945, doi:10.1371/journal.pone.0145945 (2016).

45 Zhang, L. F. et al. The Vi Capsular Polysaccharide of Salmonella Typhi Promotes Macrophage Phagocytosis by Binding the Human C-Type Lectin DC-SIGN. mBio 13, e0273322, doi:10.1128/mbio.02733-22 (2022).

46 Zhang, S. et al. SeqSero2: Rapid and Improved Salmonella Serotype Determination Using Whole-Genome Sequencing Data. Appl Environ Microbiol 85, e01746–01719, doi:10.1128/aem.01746-19 (2019).

47 Ahn, C. et al. Mechanisms of typhoid toxin neutralization by antibodies targeting glycan receptor binding and nuclease subunits. iScience 24, 102454 (2021).

48 Arricau, N. et al. The RcsB-RcsC regulatory system of Salmonella typhi differentially modulates the expression of invasion proteins, flagellin and Vi antigen in response to osmolarity. Mol Microbiol 29, 835–850, doi:10.1046/j.1365-2958.1998.00976.x (1998).

49 Liston, S. D. et al. Periplasmic depolymerase provides insight into ABC transporter-dependent secretion of bacterial capsular polysaccharides. Proc Natl Acad Sci U S A 115, E4870–e4879, doi:10.1073/pnas.1801336115 (2018).

50 Baker, S. et al. Detection of Vi-negative Salmonella enterica serovar typhi in the peripheral blood of patients with typhoid fever in the Faisalabad region of Pakistan. J Clin Microbiol 43, 4418–4425, doi:10.1128/jcm.43.9.4418-4425.2005 (2005).

51 Liaquat, S. et al. Virulotyping of Salmonella enterica serovar Typhi isolates from Pakistan: Absence of complete SPI-10 in Vi negative isolates. PLoS Negl Trop Dis 12, e0006839, doi:10.1371/journal.pntd.0006839 (2018).

52 Lee, G. Y. & Song, J. Complete genome sequence of Salmonella enterica serovar Typhi strain ISP2825. Microbiology Resource Announcements 30, 254–267 (2022).

53 Santander, J., Roland, K. L. & Curtiss, R., 3rd. Regulation of Vi capsular polysaccharide synthesis in Salmonella enterica serotype Typhi. J Infect Dev Ctries 2, 412–420 (2008).

54 Jumper, J. et al. Highly accurate protein structure prediction with AlphaFold. Nature 596, 583–589 (2021).

55 Wang, J. et al. Scaffolding protein functional sites using deep learning. Science 377, 387–394, doi:10.1126/science.abn2100 (2022).

56 Tipton, K. A. & Rather, P. N. Extraction and Visualization of Capsular Polysaccharide from Acinetobacter baumannii. Methods Mol Biol 1946, 227–231, doi:10.1007/978-1-4939-9118-1_21 (2019).

57 Hedlund, M. et al. N-glycolylneuraminic acid deficiency in mice: implications for human biology and evolution. Mol Cell Biol 27, 4340–4346 (2007).

58 Chou, H. H. et al. Inactivation of CMP-N-acetylneuraminic acid hydroxylase occurred prior to brain expansion during human evolution. Proc Natl Acad Sci U S A 99, 11736–11741, doi:10.1073/pnas.182257399 (2002).

59 Lee, S. et al. Salmonella Typhoid Toxin PltB Subunit and Its Non-typhoidal Salmonella Ortholog Confer Differential Host Adaptation and Virulence. Cell Host Microbe 27, 937–949, doi:10.1016/j.chom.2020.04.005 (2020).

60 Yang, Y. A. et al. In vivo tropism of Salmonella Typhi toxin to cells expressing a multiantennal glycan receptor. Nat Microbiol 3, 155–163, doi:10.1038/s41564-017-0076-4 (2018).

61 Saha, S. et al. Exploring the Impact of Ketodeoxynonulosonic Acid in Host-Pathogen Interactions Using Uptake and Surface Display by Nontypeable Haemophilus influenzae. mBio 12, doi:10.1128/mBio.03226-20 (2021).

62 Landig, C. S. et al. Evolution of the exclusively human pathogen Neisseria gonorrhoeae: Human-specific engagement of immunoregulatory Siglecs. Evol Appl 12, 337–349, doi:10.1111/eva.12744 (2019).

63 Deng, L. et al. Host adaptation of a bacterial toxin from the human pathogen salmonella typhi. Cell 159, 1290–1299, doi:10.1016/j.cell.2014.10.057 (2014).

64 Varki, A. & Gagneux, P. Multifarious roles of sialic acids in immunity. Ann N Y Acad Sci 1253, 16–36 (2012).

65 Thanh Duy, P., et al. Gallbladder carriage generates genetic variation and genome degradation in Salmonella Typhi. PLoS Pathog 16, e1008998, doi:10.1371/journal.ppat.1008998 (2020).

66 Keestra-Gounder, A. M., Tsolis, R. M. & Bäumler, A. J. Now you see me, now you don’t: the interaction of Salmonella with innate immune receptors. Nat Rev Microbiol 13, 206–216, doi:10.1038/nrmicro3428 (2015).

67 Siu, L. K., Yeh, K. M., Lin, J. C., Fung, C. P. & Chang, F. Y. Klebsiella pneumoniae liver abscess: a new invasive syndrome. Lancet Infect Dis 12, 881–887, doi:10.1016/s1473-3099(12)70205-0 (2012).

68 Brown, J. S. et al. The classical pathway is the dominant complement pathway required for innate immunity to Streptococcus pneumoniae infection in mice. Proc Natl Acad Sci U S A 99, 16969–16974, doi:10.1073/pnas.012669199 (2002).

69 Durmort, C. et al. Deletion of the Zinc Transporter Lipoprotein AdcAII Causes Hyperencapsulation of Streptococcus pneumoniae Associated with Distinct Alleles of the Type I Restriction-Modification System. mBio 11, e00445–00420, doi:10.1128/mBio.00445-20 (2020).

70 Roshini, J., Patro, L. P. P., Sundaresan, S. & Rathinavelan, T. Structural diversity among Acinetobacter baumannii K-antigens and its implication in the in silico serotyping. Front Microbiol 14, 1191542, doi:10.3389/fmicb.2023.1191542 (2023).

71 Chin, C. Y. et al. A high-frequency phenotypic switch links bacterial virulence and environmental survival in Acinetobacter baumannii. Nat Microbiol 3, 563–569, doi:10.1038/s41564-018-0151-5 (2018).

72 Holt, K. E. et al. Genomic analysis of diversity, population structure, virulence, and antimicrobial resistance in Klebsiella pneumoniae, an urgent threat to public health. Proc Natl Acad Sci U S A 112, E3574–3581, doi:10.1073/pnas.1501049112 (2015).

73 Wyres, K. L., Lam, M. M. C. & Holt, K. E. Population genomics of Klebsiella pneumoniae. Nat Rev Microbiol 18, 344–359, doi:10.1038/s41579-019-0315-1 (2020).

74 Pan, Y. J. et al. Genetic analysis of capsular polysaccharide synthesis gene clusters in 79 capsular types of Klebsiella spp. Sci Rep 5, 15573, doi:10.1038/srep15573 (2015).

75 Soni, P. N., Hoosen, A. A. & Pillay, D. G. Hepatic abscess caused by Salmonella typhi. A case report and review of the literature. Dig Dis Sci 39, 1694–1696, doi:10.1007/bf02087778 (1994).

76 Ciraj, A. M., Reetika, D., Bhat, G. K., Pai, C. G. & Shivananda, P. G. Hepatic abscess caused by Salmonella typhi. J Assoc Physicians India 49, 1021–1022 (2001).

77 Gömceli, I., Gürer, A., Ozdoğan, M., Ozlem, N. & Aydin, R. Salmonella typhi abscess as a late complication of simple cyst of the liver: a case report. Turk J Gastroenterol 17, 151–152 (2006).

78 Giorgio, A., Tarantino, L. & De Stefano, G. Hepatic abscess caused by Salmonella typhi: diagnosis and management by percutaneous echo-guided needle aspiration. Ital J Gastroenterol 28, 31–33 (1996).

79 Jorge, J. F. et al. Salmonella typhi liver abscess overlying a metastatic melanoma. Am J Trop Med Hyg 90, 716–718, doi:10.4269/ajtmh.13-0573 (2014).

80 Chaudhry, R. et al. Unusual presentation of enteric fever: three cases of splenic and liver abscesses due to Salmonella typhi and Salmonella paratyphi A. Trop Gastroenterol 24, 198–199 (2003).

81 Chang, S. J., Song, J. & Galan, J. E. Receptor-Mediated Sorting of Typhoid Toxin during Its Export from Salmonella Typhi-Infected Cells. Cell Host Microbe 20, 682–689, doi:10.1016/j.chom.2016.10.005 (2016).

82 Shen, W., Le, S., Li, Y. & Hu, F. SeqKit: A Cross-Platform and Ultrafast Toolkit for FASTA/Q File Manipulation. PLoS One 11, e0163962, doi:10.1371/journal.pone.0163962 (2016).

83 Varki, N. M., Strobert, E., Dick, E. J., Jr., Benirschke, K. & Varki, A. Biomedical differences between human and nonhuman hominids: potential roles for uniquely human aspects of sialic acid biology. Annu Rev Pathol 6, 365–393, doi:10.1146/annurev-pathol-011110-130315 (2011).

84 Neupane, D. P., Ahn, C., Yang, Y. A., Lee, G. Y. & Song, J. Malnutrition and maternal vaccination against typhoid toxin. PLoS Pathog 18, e1010731, doi:10.1371/journal.ppat.1010731 (2022).

85 Dulal, H. P. et al. Neutralization of typhoid toxin by alpaca-derived, single-domain antibodies targeting the PltB and CdtB subunits. Infect Immun 90, e0051521, doi:10.1128/iai.00515-21 (2022).

